# Adrenergic signaling induces a pro-tumorigenic B cell state in colorectal cancer

**DOI:** 10.64898/2026.01.23.700260

**Authors:** Meike S. Thijssen, Simone L. Schonkeren, Linde Coolkens, Yuchi Zou, Lieve Temmerman, Joëlle de Vaan, Musa Idris, Jorunn Vrancken, Kim Wouters, Nathalie Vaes, Marion J. Gijbels, Sharon Scardellato, Erwin Wijnands, Willine van de Wetering, Frank Verhaegen, Ludwig J. Dubois, Erik Biessen, Carmen López Iglesias, Kim M. Smits, Alberto Bardelli, Stefano Casola, Werend Boesmans, Veerle Melotte

## Abstract

The importance of neuron-tumor crosstalk has gained increasing attention, yet its influence on the cellular and molecular landscape of colorectal cancer (CRC) remains largely unexplored. Here, we show that colonic innervation shapes the tumor immune microenvironment in a murine model of colitis-associated CRC. Although neuronal density does not affect tumor number, size, or overall burden, transcriptomic profiling of cells isolated from the tumor of hypo-innervated mice revealed extensive differential gene expression, including genes involved in immunoglobulin (Ig) signaling and the cancer-relevant hallmark ‘avoiding immune destruction’. Flow cytometry analysis of leukocyte populations demonstrated a significant reduction of B cells in the cancerous colon of hypo-innervated mice, notably a decrease in germinal center B cells and an altered class-switching profile, characterized by reduced IgA and increased IgD expression. Fluorescence and transmission electron microscopy showed that colonic B cells are primarily localized in the submucosa near neuronal processes containing varicose release sites, suggesting direct neuron-B cell interactions. Functional assays revealed that adrenergic stimulation of B cells promotes their proliferation and maturation, enhances IL-10 secretion, and alters immunoglobulin profiles. Strikingly, the transcriptome of epinephrine-treated B cells closely mirrors that of plasma cells from human CRC tissues, and the transcriptomic signature of these epinephrine-stimulated B cells associates with poorer patient survival. Together, these findings uncover an adrenergic neuron - B cell axis in CRC, providing evidence for direct neuroimmune interactions that affect B cell maturation and may influence tumor progression and therapeutic responses.

## Introduction

Nerves, as active members of the tumor microenvironment (TME)^1,2^, have been recognized to influence tumor biology and their abundance correlates with patient outcomes^3,4^. The peripheral nervous system innervates nearly all organs and is therefore in a prime position to provide nerve supply to the TME in multiple cancer types^5^. Among visceral organs, the gastrointestinal (GI) tract is particularly densely innervated. Next to extrinsic sympathetic and parasympathetic nerves, it harbors an intrinsic enteric nervous system (ENS)^6^, positioning nerves to significantly influence colorectal cancer (CRC) development and progression^7–9^. However, few studies have addressed how neurons shape the tumor and its environment in colorectal cancer (CRC)^10,11^.

Early observations linked CRC to perineural invasion^12–14^, axonogenesis^15^, and neoneurogenesis^16^, each associated with tumor progression and poor prognosis. Yet, experimental models manipulating innervation have produced inconsistent results^17–20^. This may reflect the diverse neuronal subtypes and signaling pathways involved. For instance, our recent work showed that vasoactive intestinal peptide (VIP)- and neuronal nitric oxide synthase (nNOS)-expressing fibers directly innervate the CRC stroma, while other subtypes like cholinergic or serotonergic nerves remain peritumoral^21^.

Beyond direct interactions, indirect communication routes between neurons and tumor cells in CRC are complex and have only just began to be uncovered. We previously demonstrated that neurons can influence CRC indirectly by modulation of the extracellular matrix (ECM), as the neuron-derived ECM proteins Fibulin-2 and Nidogen-1, regulated by N-Myc Downstream-Regulated Gene 4 (NDRG4), significantly affect intestinal tumor growth^22,23^. In addition, adrenergic neuronal stimulation has been found to induce secretion of nerve growth factor by cancer-associated fibroblasts, which promotes CRC growth^24^. Recent studies also indicate that neuronal signals can modulate immune cell infiltration and function in multiple cancers^25,26^. For example, sympathetic signaling was shown to enable macrophage infiltration and to induce a pro-tumorigenic phenotype switch^27^, as well as hampering T cell activation and anti-tumor responses^28,29^. Nevertheless, despite growing insights, the spatial relations and signaling modules of neuro-immune interactions in CRC remain poorly defined.

To systematically assess how alterations in neuronal innervation impact the CRC microenvironment in an unbiased way, we made use of a transgenic mouse model with reduced intestinal innervation (hypo-innervated). We demonstrate that decreased colonic innervation significantly reshapes the immune landscape of colitis-associated CRC tumors. Hypo-innervation specifically impacts B cell abundance and maturation, highlighted by reductions in germinal center B cells and shifts in immunoglobulin expression profiles. Furthermore, we identified direct neuron–B cell interactions, supported by the proximity of colonic B cells to neurotransmitter release sites. Adrenergic stimulation of B cells recapitulates molecular signatures observed in human CRC plasma cells, and this epinephrine-treated B cell signature is associated with poorer patient survival.

## Results

### Colonic innervation density does not alter tumor development, but affects tumor transcriptome

*Hand2^fl/+^;Wnt1-Cre2* mice, which exhibit reduced innervation density due to conditional *Hand2* deletion in neural crest-derived cells^30,31^, and control *Hand2^fl/+^*mice were used to assess the impact of neuron density on CRC. *Hand2^fl/+^;Wnt1-Cre2* mice had no significant differences in body weight (Supplementary Figure 1A-B) or colon length (Supplementary Figure 1C) compared to *Hand2^fl/+^* control littermates before induction of colitis-associated cancer. Myenteric neuron numbers were significantly reduced in both the proximal (Supplementary Figure 1D; 528±94 neurons/mm² and 431±82 neurons/mm²) and distal (Supplementary Figure 1E; 266±71 neurons/mm² and 203±51 neurons/mm²) colon of *Hand2^fl/+^;Wnt1-Cre2* mice evaluated by antibody labeling for the pan-neuronal marker Hu (Supplementary Figure 1F), confirming hypo-innervation in the colon.

*Hand2^fl/+^;Wnt1-Cre2* and *Hand2^fl/+^*mice (both males and females) were subjected to an AOM/DSS CRC induction protocol and tumor burden was tracked *in vivo* by micro CB-CT scans throughout the protocol to assess tumor development (Figure 1A). Tumor initiation and tumor burden, i.e. the total tumor volume per mouse, did not significantly differ between *Hand2^fl/+^* and *Hand2^fl/+^;Wnt1-Cre2* mice during the protocol (Figure 1B). At end-stage, colons from control CRC mice had decreased in length compared to colons from naïve mice (*P*=0.007), indicative of severe DSS-induced colitis, but did not shorten significantly in *Hand2^fl/+^;Wnt1-Cre2* mice (*P*=0.675) (Figure 1C and Supplementary Figure 1C). Tumor number, tumor size and tumor burden did not significantly differ between *Hand2^fl/+^;Wnt1-Cre2* and *Hand2^fl/+^* mice (Figure 1D-F). Furthermore, histological tumor classification was not found to be different between both genotypes (Figure 1G-H).

**Figure 1.**
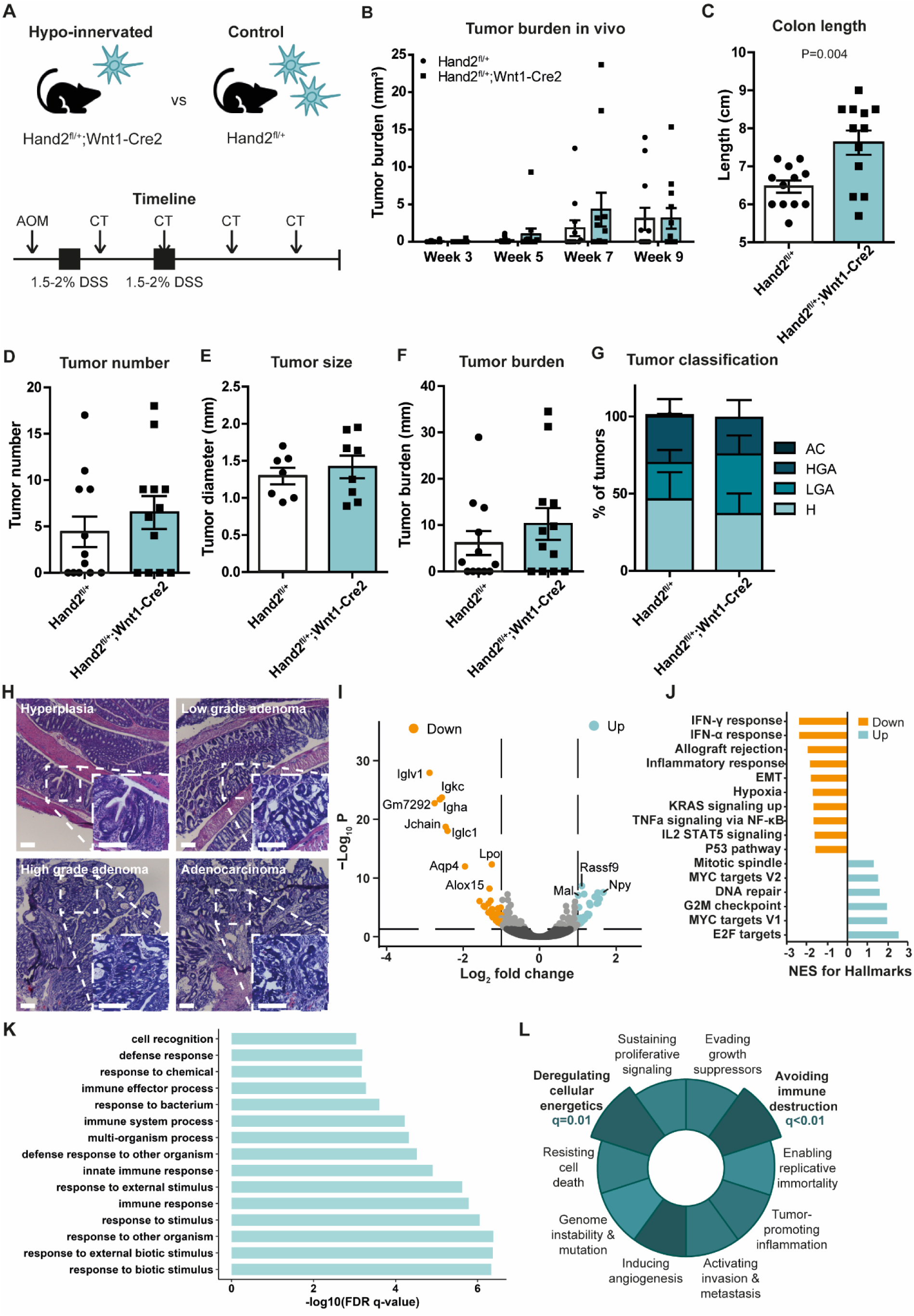
| Colon and tumor characteristics during and after AOM/DSS. **A|** Timeline for AOM/DSS procedure to induce CRC, including timepoints at which CT scans were acquired. **B|** *In vivo* tumor tracking with CT scans shows no significant difference between *Hand2^fl/+^;Wnt1-Cre2* (N=12) and *Hand2^fl/+^* (N=12) mice in tumor burden. Tumor burden was calculated by summing all tumor volumes (mm³) per mouse. **C|** *Hand2^fl/+^* mice have significantly shorter colons than *Hand2^fl/+^;Wnt1-Cre2* mice. **D|** Tumor number, **E|** size (averaged per mouse, N=7 and N=8, mice without tumors were excluded from analysis) and **F|** burden (tumor number multiplied by average tumor size) are not significantly different between *Hand2^fl/+^;Wnt1-Cre2* and *Hand2^fl/+^* mice. **G|** There is no significant difference in relative frequency of the different tumor classifications between *Hand2^fl/+^* and *Hand2^fl/+^;Wnt1-Cre2* mice. **H|** Colon tumors were classified as hyperplasia (H), low grade adenoma (LGA), high grade adenoma (HGA), or adenocarcinoma (AC) based on histology. Scale bar is 100 µm. **I|** Volcano plot showing the mean log_2_-transformed fold change (*x*-axis) and −log_10_-transformed *P*-value of differentially expressed genes between tumors from *Hand2^fl/+^* (N=3) and *Hand2^fl/+^;Wnt1-Cre2* (N=2) mice. Dashed lines indicate the threshold of significant gene expression, defined as log_2_-transformed fold change ≤−1 and ≥1 with −log_10_(P)≥1.301. **J|** The top 10 significantly decreased gene sets and all six significantly increased gene sets are shown with the normalized enrichment score. **K|** The top 15 significant biological pathways (-log_10_-transformed *Q*-value) from gene ontology analysis include ‘(defense) response to bacterium’ and ‘innate immune response’. **L|** GSEA was performed on hallmarks of cancer gene sets. The hallmarks ‘deregulating cellular energetics’ and ‘avoiding immune destruction’ were significantly enriched. All data are presented as mean ±SEM.

However, tumors of *Hand2^fl/+^;Wnt1-Cre2* and *Hand2^fl/+^* mice differ significantly at the transcriptomic level. RNA sequencing analysis revealed that a reduced colonic innervation affects the tumor phenotype by a downregulation of, among others, immunoglobulin (Ig)-related transcripts (e.g. *Igha*, *Igkc*, *Iglc1, Iglv1* and *Jchain*; Figure 1I). Gene set enrichment analysis (GSEA) showed that downregulated gene sets in tumors isolated from *Hand2^fl/+^;Wnt1-Cre2* mice relate to immune signaling, including interferon-γ (IFN-γ) and interferon-α (IFN- α) response, allograft rejection, inflammatory response, TNF-α signaling, and IL2 STAT5 signaling (Figure 1J). Furthermore, among the upregulated gene sets are DNA repair and G2M checkpoint, while downregulated gene sets include p53 pathway and epithelial-mesenchymal transition, indicating that colonic hypo-innervation results in downregulation of pro-tumorigenic pathways and upregulation of anti-tumorigenic pathways. Gene ontology analysis defined that the differentially expressed genes relate to the biological pathways *response to other organisms* and *external (biotic) stimulus* as well as *immune response*, which include the pathways (*defense) response to bacterium* and *innate immune response* (Figure 1K). In addition, a cancer hallmark GSEA showed that differentially expressed genes are predominantly enriched in the two hallmarks ‘deregulating cellular energetics’ and ‘avoiding immune destruction’ (Figure 1L)^32,33^.

### Reduced colonic innervation has no effect on intestinal microbiota composition and tumor cell metabolism

Since the transcriptomic analysis shows enrichment of bacterium-related gene ontology terms, we investigated whether alterations in intestinal microbiota could contribute to the observed tumor transcriptome changes in the hypo-innervation model. In line with previous findings^34–36^, the fecal microbiome was significantly affected by AOM/DSS treatment (beta-diversity weighted UniFrac *P*=0.008), showing enrichment of the bacterial families *Bacteroidaceae* and *Akkermansiaceae* (0.00%±0.00% before and 12.94%±9.47% after AOM/DSS, *W*=38; and 0.04%±0.08% before and 5.39%±3.82% after AOM/DSS, *W*=32, respectively) (Supplementary Figure 2A-B). However, the reduced colonic neuronal density did not affect the intestinal microbiome, neither in healthy conditions (Supplementary Figure 2C, E) nor after cancer induction (Supplementary Figure 2D, F).

Directed by the enrichment of differentially expressed genes belonging to the cancer hallmark ‘deregulating cellular energetics’, SCENITH was employed to study the metabolism of tumor cells of *Hand2^fl/+^;Wnt1-Cre2* and *Hand2^fl/+^* mice at the single cell level (Supplementary Figure 3A-C). As suggested by the literature^37^, for energy supply colonic tumor cells were found to be much more dependent on glycolysis than on oxidative phosphorylation; however, no differences in these main metabolic pathways were observed between tumor cells isolated from hypo-innervated and control littermates (Supplementary Figure 3D-E).

### Germinal center B cells are reduced and the immunoglobulin profile is altered in the cancerous colon of hypo-innervated mice

The enrichment of differentially expressed genes involved in immune responses and immune signaling prompted us to determine the abundance of the major lymphoid and myeloid immune cell populations using flow cytometry in the AOM/DSS-treated mouse colon^38^ (gating strategies depicted in Supplementary Figure 4A-B). After the induction of CRC, colonic B cell abundance was significantly reduced in *Hand2^fl/+^;Wnt1-Cre2* as compared to *Hand2^fl/+^* mice (Figure 2A, C), while myeloid populations were found to be unaltered (Figure 2B). Further characterization of B cell subtypes (Supplementary Figure 4C) revealed a significant decrease in GL7^+^ germinal center (GC) B cells in the cancerous colon of hypo-innervated mice (Figure 2D-E). Furthermore, immature B cells were significantly decreased (Figure 2D), as suggested by an overrepresentation of IgD-expressing mature B cells, whereas IgM expression was unaltered (Figure 2F). Surface IgA-expressing B cells were reduced in frequency in the cancerous colon of *Hand2^fl/+^;Wnt1-Cre2* mice when compared to controls (Figure 2F). The same differences were neither observed in healthy (Figure 2G-I), nor in DSS-treated colitis transgenic animals when compared to control mice (Figure 2J-L), concluding that the decrease in colonic GC and IgA-expressing B cells in hypo-innervated mice is instructed by the CRC environment. B cell subset analysis of mesenteric lymph nodes was also performed, excluding significant differences between control and *Hand2* haploinsufficient mice, therefore indicating that the colonic B cell phenotype of hypo-innervated mice is independent from the systemic B cell immunity triggered in mesenteric lymph nodes (Supplementary Figure 5A). As the percentage of immune cells present in the mouse colon steadily increased from healthy (*Hand2^fl/+^*=0.6% of live cells, *Hand2^fl/+^;Wnt1-Cre2*=0.9% of live cells) to inflamed (*Hand2^fl/+^*=7.2% of live cells, *Hand2^fl/+^;Wnt1-Cre2*=7.9% of live cells) to cancerous colon (*Hand2^fl/+^*=19.3% of live cells, *Hand2^fl/+^;Wnt1-Cre2*=25.1% of live cells) (Supplementary Figure 5B), so did the total fraction of B cells and that of GC B cells in control mice (Supplementary Figure 5C-D). In contrast, hypo-innervated mice showed a blunted increase in GC B cells from the inflamed to the cancerous stage (Supplementary Figure 5C-D). This provides support for the idea that the B cell response, in particular that of GC B cells, to colonic carcinogenesis critically depends on optimal innervation of the colonic tissue. Specifically, these results suggest that colonic innervation regulates B cell class-switching in GCs and maturation towards Ig-producing plasma cells in CRC.

**Figure 2.**
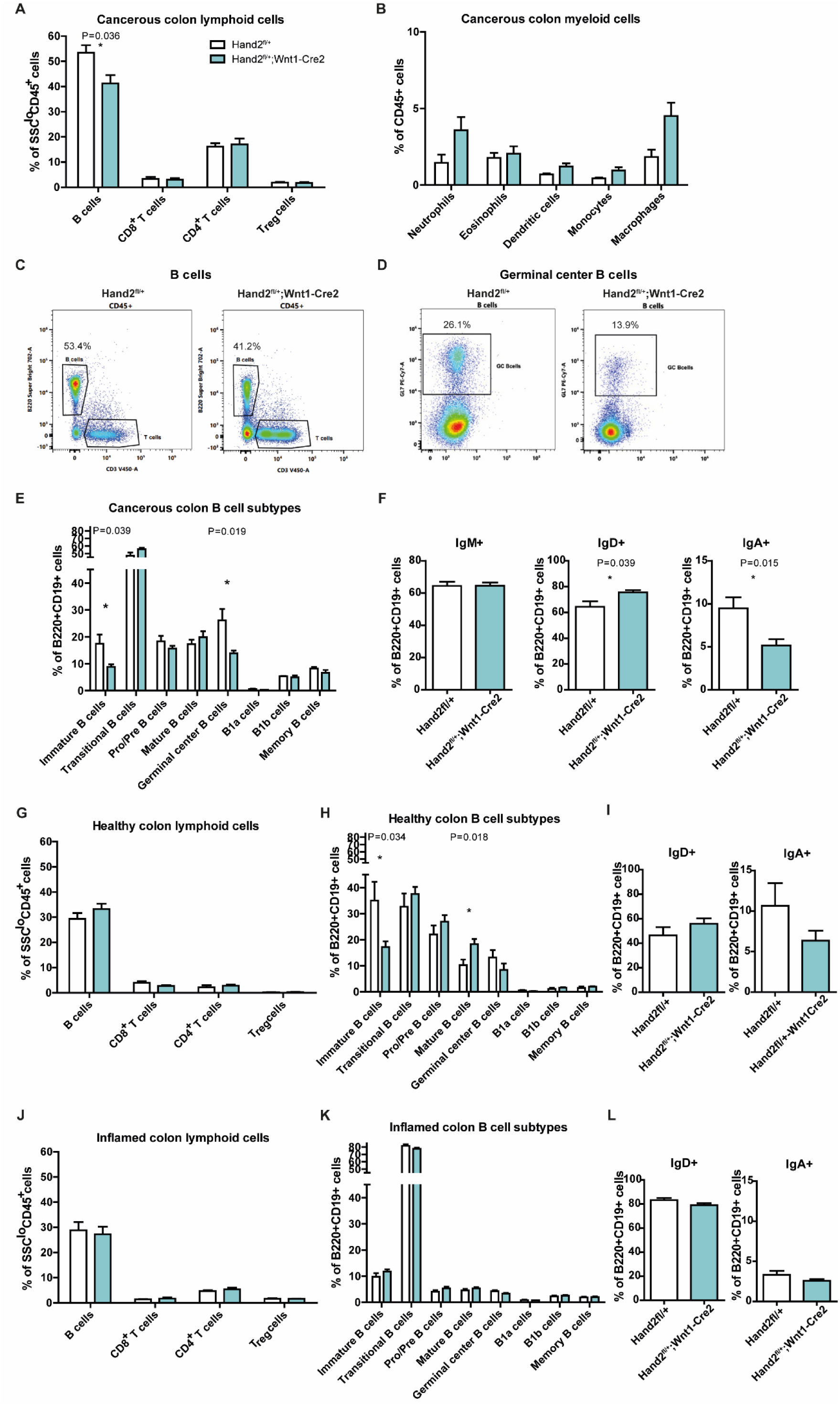
| Flow cytometry analysis of immune cell populations in the healthy, inflamed and cancerous colon of *Hand2^fl/+^* and *Hand2^fl/+^;Wnt1-Cre2* mice. **A|** Analysis of lymphoid cells in the cancerous colon shows a significant reduction in the number of B cells in *Hand2^fl/+^;Wnt1-Cre2* (N=7) compared to *Hand2^fl/+^* (N=4) mice. **B|** Myeloid cells are not affected by hypo-innervation. **C|** Flow cytometry plots of B cells **D|** and germinal center (GC) B cells depict the decrease in population abundance in *Hand2^fl/+^;Wnt1-Cre2* mice. **E|** B cell subtype analysis identifies the specific downregulation of GC B cells and immature B cells in hypo-innervated cancerous colon (N=8) compared to control cancerous colon (N=8). **F|** Immunoglobulin expression on B cells is altered by an increase in IgD^+^ and a decrease in IgA^+^ B cells. **G|** In healthy colon, no differences in lymphoid populations were observed in *Hand2^fl/+^;Wnt1-Cre2* (N=8) compared to *Hand2^fl/+^* (N=8) mice. **H|** B cell subtype analysis shows a shift towards more mature B cells in hypo-innervated colon, but GC B cells are not affected. **I|** Relative abundance of immunoglobulin-expressing B cells remains unaltered. In inflamed colons (N=8 vs 8), **J|** lymphoid populations, **K|** B cell subtypes, and **L|** Ig-expressing B cells are not affected by a decreased neuronal density. All data are presented as mean ±SEM.

### Colonic B cells are in a prime position and state to respond to neuronal signals

To explore the spatial context in which neurons and B cells interact, we examined the localization of B cells relative to nerve fibers in the colon of AOM/DSS-treated mice. B cells are mostly present in the submucosal layer of the colon wall either as single cells or organized in isolated lymphoid follicles, but they could also be found in the tumor stroma (Figure 3A-C). Consistent with their submucosal localization, colonic B cells were frequently found in close proximity to nerve fibers, also when clustered in lymphoid follicles (Figure 3A-B). Interestingly, despite the sparse innervation of tumor tissue^21^, intratumoral B cells are similarly located in the vicinity of nerve fibers (Figure 3C). The close spatial association between neuronal processes and B cells was confirmed by electron microscopy of human CRC patient tissues (Figure 3D), which further demonstrated that neurotransmitter-vesicle-containing processes, only partially enveloped by enteric glia, neighbor B cells. Given the varicose release nature of intestinal innervation provided by the autonomic nervous system, these data suggest that B cells are in a prime position to receive neurochemical input.

**Figure 3.**
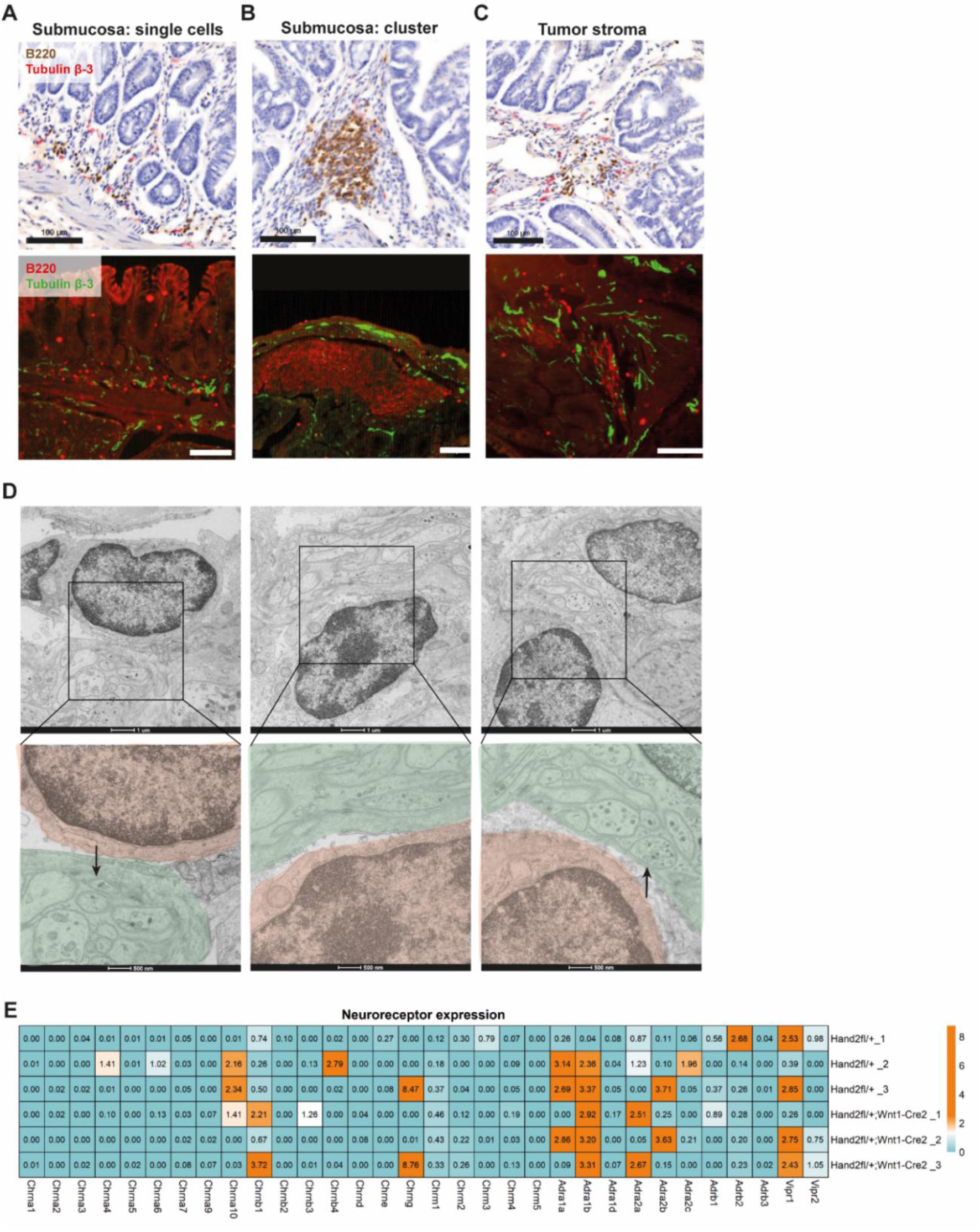
| B cell – neuron interactions in mouse and human CRC. **A|** B cells (↑brown, ↓red) are located in close proximity to neuronal processes (↑red, ↓green) in the submucosal layer of the colon as single cells, or **B|** as lymphoid clusters. **C|** In AOM/DSS-treated *Hand2^fl/+^;Wnt1-Cre2* mice, B cells are also present in the tumor stroma in vicinity of nerve fibers. Scale bars are 100 µm. **D|** Electron microscopy imaging confirms the close proximity of nerve fibers (green) and B cells (red) in human CRC tissues, and neurotransmitter vesicles are observed in fibers neighboring B cells (arrows). Scale bars are 1 µm and 500 nm. **E|** The colonic B cells of *Hand2^fl/+^*and *Hand2^fl/+^;Wnt1-Cre2* mice express cholinergic (Chrn/Chrm), adrenergic (Adr) and VIPergic (Vipr) receptor family members (N=6), with no effect of innervation level on neurotransmitter receptor profile.

To identify possible routes of neurochemical communication between neurons and B cells, we performed bulk RNA sequencing of B cells isolated from colons of AOM/DSS-treated *Hand2^fl/+^;Wnt1-Cre2* and *Hand2^fl/+^* mice. We found that colonic B cells express a repertoire of neuroreceptors, including cholinergic (Chrn and Chrm), adrenergic (Adr) and VIPergic (Vipr) receptors (Figure 3E). Taken together, these results suggest that neurons can directly communicate with B cells in the cancerous colon, possibly via neurotransmitters such as acetylcholine, epinephrine and/or VIP.

### Epinephrine induces the neuronal phenotype characterized by increased maturation in B lymphocytes

To investigate whether acetylcholine, epinephrine and/or VIP influence B cell behavior, as suggested by the results from our hypo-innervated mouse model, we monitored the effect of these neurotransmitters in an *in vitro* culture model of human GC B cells isolated from the tonsil of healthy volunteers (Figure 4A). Primary GC B cells were cultured for 10 days on a feeder layer of immortalized follicular dendritic cell Y46 cells engineered to constitutively express CD40L and IL21^39^. Addition of epinephrine to the culture medium, but not acetylcholine and VIP, induced a significant increase in the number of viable cells over a four-day culture period (Figure 4B), suggesting that the reduced fraction of B cells observed in hypo-innervated mice is, at least in part, induced by hampered proliferation.

**Figure 4.**
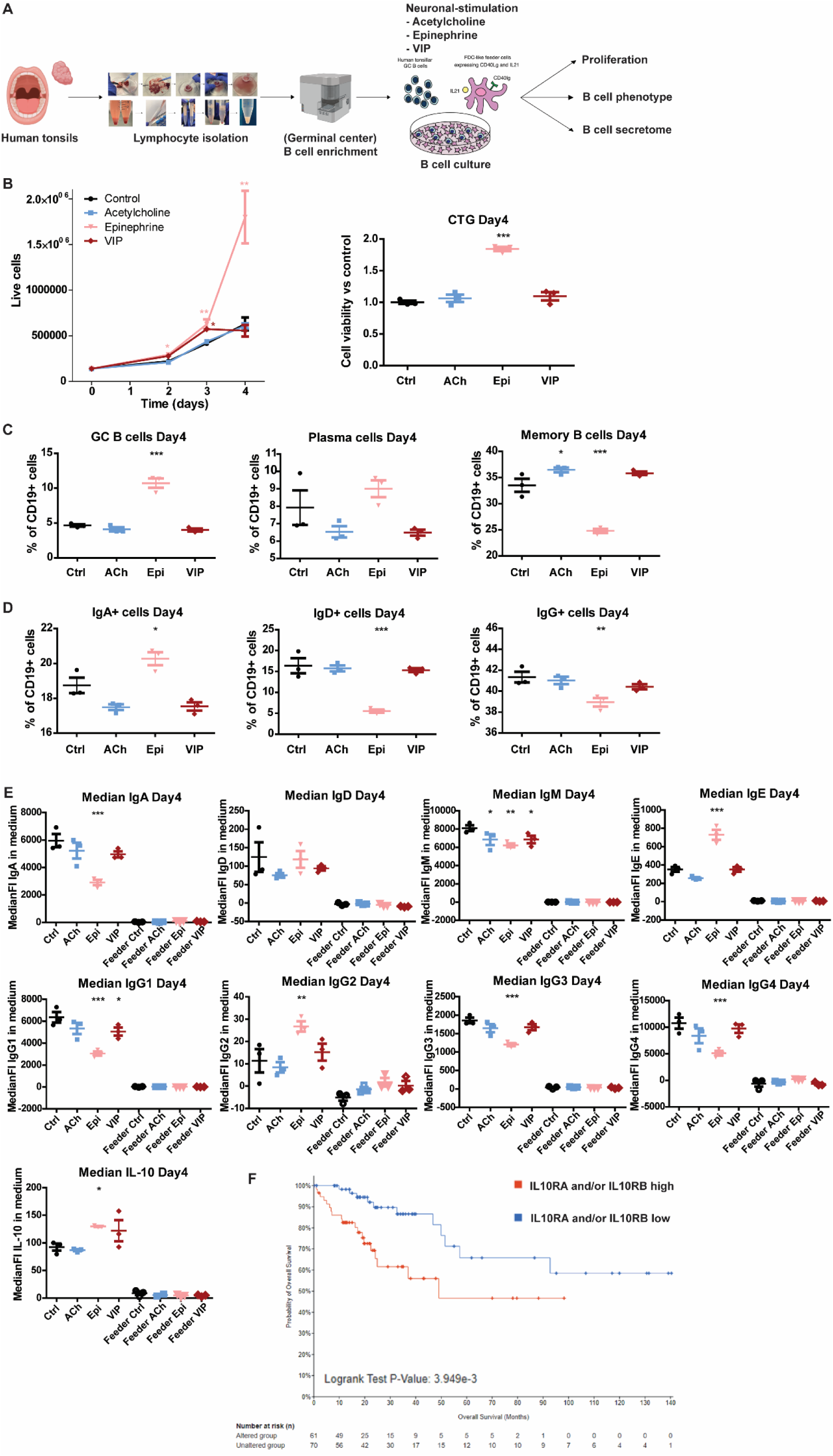
| The response of human primary B cell cultures to neurotransmitter stimulation. **A|** Human primary B cell cultures were made from healthy tonsils, enriched for GC B cells, cultured on feeder cells and stimulated with the neurotransmitter acetylcholine (Ach, blue; 200 μM), epinephrine (Epi, pink; 100 μM) or VIP (red; 500 nM). **B|** Epinephrine increased the amount of live B cells, and B cell viability by CellTiter-Glo luminescent assay (N=3). **C|** Flow cytometric analysis shows an increase in CD38^+^IgD^-^CD27^+^CD10^+^ GC B cells and a decrease in CD38^-^IgD^-^CD27^+^CD10^+^ memory B cells after stimulation with epinephrine for 4 days. **D|** Epinephrine boosts the abundance of IgA^+^ B cells while IgD^+^ and IgG^+^ B cells are reduced. **E|** Secretome analysis indicates an effect of epinephrine on immunoglobulin and IL-10 secretion from B cells. **F|** IL-10 receptor (IL10RA/IL10RB) expression in human CRC cases from TCGA is associated with poor prognosis. All data are presented as mean ±SEM.

Next, we determined whether neurotransmitter stimulation influenced the B cell phenotype. Flow cytometry analyses (Supplementary Figure 6A) revealed an epinephrine-induced enrichment for cells presenting a GC B cell phenotype (CD38^+^IgD^-^CD27^+^CD10^+^), while those presenting a post-GC memory B cell phenotype (CD38^-^IgD^-^CD27^+^CD10^+^) are reduced in frequency when compared to unstimulated cultures after four days (Figure 4C). This phenotypic switch persisted for up to ten days of culture. CD38^+^IgD^-^CD27^+^CD10^-^ plasma cells showed a significant increase in percentage over their unstimulated counterparts at later time points (between seven to ten days of stimulation) (Supplementary Figure 6B-D). Notably, the abundance of IgA^+^ B cells increased while IgD^+^ and IgG^+^ B cell subsets decreased in frequency after exposure to epinephrine (Figure 4D). While sympathetic efferents have been shown to receive input from enteric neurons to impact adrenergic neuronal activity^40,41^, we also observed a reduction in tyrosine hydroxylase (TH) immunoreactivity, as a marker for sympathetic innervation, in the colonic myenteric plexus of *Hand2^fl/+^;Wnt1-Cre2* mice (Supplementary Figure 7A-B). This indicates that next to indirect modulation by the ENS, impaired adrenergic signalling could also directly contribute to the *in vivo* phenotype. Merging *in vivo* mouse data with *in vitro* human GC B cell culture studies, we propose that reduced adrenergic signalling is responsible for the underrepresentation of total B cells and in particular of the GC and surface IgA+ fraction in the hypo-innervated cancerous colon.

Secretome analysis using a B cell immunoglobulin and cytokine panel showed that the immunoglobulin secretion from *in vitro* cultured human GC B cells was altered by administration of epinephrine to the culture medium. Specifically, we observed decreased soluble IgA, IgM, IgG1, IgG3 and IgG4 titers, while IgG2 and IgE increased in the culture medium (Figure 4E). Moreover, epinephrine augmented the release of the anti-inflammatory cytokine IL-10 (Figure 4E). The Cancer Genome Atlas (TCGA) analysis further showed that high IL-10 receptor expression in human CRC is associated with poor prognosis, possibly suggesting that increased IL-10 release and signaling from epinephrine-treated B cells is unfavorable (Figure 4F).

### The adrenergic neuron-B cell communication axis is pro-tumorigenic

To validate the functional relevance of neuronal modulation of B cells in CRC, we performed transcriptomic profiling of primary human GC B cells stimulated with neuronal cues. Among the tested neurotransmitters, epinephrine induced the most pronounced transcriptional changes (Figure 5A), with differentially expressed genes enriched in pathways and molecular functions related to immunoglobulin signaling, B cell activation, and cytokine-mediated immune responses (Figure 5B-D). These analyses support a model in which adrenergic signaling skews B cell function towards an activated or plasma cell-like phenotype.

**Figure 5.**
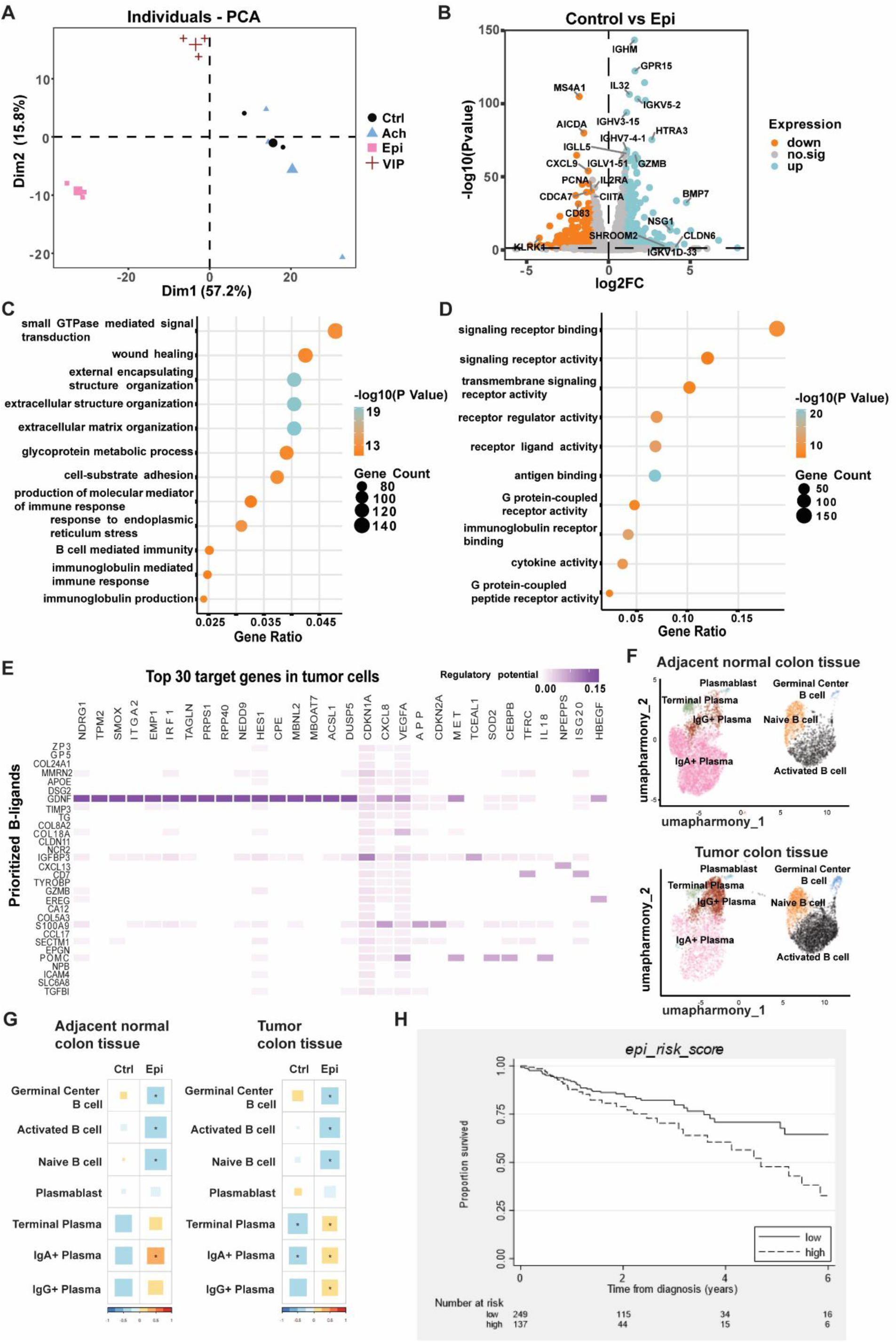
| Epinephrine - B cell communication at transcriptomic level and its links to colorectal cancer. **A|** Principal Component Analysis (PCoA) of the B cell transcriptome confirms that epinephrine has the biggest effect on B cell phenotype (N=3). **B|** Differentially expressed genes between control and epinephrine-treated B cells are enriched for immunoglobulin and immunity-related genes. The effect of epinephrine on B cell immunoglobulin and cytokine signaling is further shown by **C|** GO term analysis and **D|** GO molecular function analysis. **E|** Nichenet analysis of differential gene expression between epinephrine-treated and untreated B cells for B cell - cancer interaction showed several candidates for the shifted B cell - tumor crosstalk upon epinephrine exposure. Of note: B cell was considered as sender and CRC cell as receiver. **F|** UMAP harmony plot of the B cell and plasma cell groups of human normal colon and CRC tissues based on single cell sequencing data from Kobayashi et al. **G|** Epinephrine-treated B cells are more similar to the plasma cell populations in the tumor tissue compared to normal colon. **H|** The epinephrine-induced B cell signature (group high= dotted line) is associated with a worse survival for CRC patients as visualized with Kaplan-Meijer plot (P=0.0214).

To explore whether such neuronally reprogrammed B cells can also influence tumor cell behavior via paracrine signaling, we performed a NicheNet analysis, using the epinephrine-responsive genes as candidate ligands and highly expressed CRC genes as potential targets (Figure 5E). This analysis identified several biologically relevant ligand-target pairs. Among others, multiple collagen genes (*Col24a1, Col8a2, Col5a3* and *Col18a1*), EGFR-ligands (*Ereg* and *Epgn*), and B/T cell regulators (*Cd7, CCL17, Gzmb, Tyrobp* and *Cxcl13*) were identified as potential mediators involved in epinephrine-treated B cell to CRC cell signaling, highlighting the potential of neuronally-altered B cells in modulating the colorectal TME and carcinogenesis.

Next, we compared the resulting gene expression signature to single-cell RNA-sequencing data of colorectal tumor tissue and adjacent normal tissue, previously described by Lee et al.^42^. We found that GC B cells are increased in CRC tissues compared to healthy neighboring colon tissue, resembling the changes induced by epinephrine on B cells *in vitro* (Figure 5F). Moreover, the transcriptome of epinephrine-stimulated B cells was found to be more similar to that of tumor plasma cell populations compared to that of plasma cells of the normal colon (Figure 5G). Strikingly, the epinephrine-induced B cell signature, defined by genes significantly upregulated in treated versus untreated B cells, associates with a reduced survival in the colon adenocarcinoma (COAD)-TCGA CRC patient group (P=0.0214, Figure 5H, Supplementary Table 3), even after adjusting for sex and age (HR: 1.652, P=0.031, 95% CI: 1.048-2.605). Notably, this adverse prognostic association was not restricted to CRC: the same signature similarly correlated with poorer outcome in gastric cancer (Supplementary Figure 8), suggesting that adrenergic shaping of B-cell states may represent a broader feature of gastrointestinal tumor biology. Together, these data suggest that adrenergic neuronal signaling may skew B cell differentiation states toward immune-regulatory and tumor-supportive profiles that are observed in cancer patients and linked to worse clinical outcomes.

## Discussion

To gain insight into how neuronal input shapes the tumor microenvironment in CRC, we utilized the *Hand2^fl/+^;Wnt1-Cre2* mouse model, which exhibits reduced intestinal innervation, in combination with AOM/DSS-induced colitis-associated CRC^43^. Tumor incidence, size, and histology are not affected between hypo-innervated and control mice, but transcriptomic profiling revealed profound differences in tumor and TME biology. Notably, two hallmarks of cancer, i.e. ‘deregulating cellular energetics’ and ‘avoiding immune destruction’^44,45^, were found to be enriched for differentially expressed genes. The downregulation of several inflammation-related gene sets indicates a reduced inflammatory response in tumors of hypo-innervated mice. Remarkably, hypo-innervation negatively affects the transcript levels of immunoglobulin-related genes and the abundance of B cells in the cancerous colon, reflected by a decrease in the number of GC B cells and immature B cells, and a change in immunoglobulin expression on B cells, with a decrease in IgA and an increase in IgD-positive subsets. Together, the downregulation of immunoglobulin-associated gene expression in the tumor, the reduction of B cells in the cancerous colon of hypo-innervated mice, and the alterations of immunoglobulin heavy chain class choice on these B cells are indicative of a reduced immune activation status and inflammatory response^46,47^.

More than 80% of activated B cells in the body are located at mucosal sites, including those of the intestines, where they provide a first-line defense against pathogens^48^. Here, we show that colonic B cells are mostly located in the submucosa as single cells or lymphoid clusters as well as in the tumor stroma. Interestingly, nerve fibers innervate B cell clusters, as suggested by the close proximity of neurotransmitter varicose release sites to individual B cells, consistent with previous observations^49,50^. Multiple studies have indicated that neurotransmitters, such as acetylcholine^51^, (nor)epinephrine^52^, VIP^53^ and substance P^54^, directly influence B cell phenotype and function by binding to their corresponding receptors, at least in *in vitro* cell line settings^55^. We confirm, at both the transcriptomic and functional level, that primary colonic B cells express neurotransmitter receptors, including cholinergic, adrenergic and VIP receptors. B cell frequency is increased during colitis and AOM/DSS-induced colitis-associated cancer and B lymphocytes are implicated in the regulation of chronic intestinal inflammation^46,56^. Our experiments indicate that hypo-innervated mice suffer from an impaired recruitment of B cells, GC B cells in particular, to inflamed and established colorectal tumors, suggesting a role for the peripheral nervous system in modulating B cell responses in chronic immunogenic and cancerous states of the colon. In CRC patients, GC B cells are enriched in proximity to tumor-infiltrated colonic tissue compared to healthy colon^57^. In GCs, B cells undergo rapid clonal expansion, Ig somatic hypermutation, and are subjected to stringent antigen-driven selection. This process is accompanied by the progressive replacement of IgM^+^/IgD^+^ B cells with their IgH class-switched counterparts (i.e. IgG, IgA) as a result of preferential expansion of isotype-switched GC B cells^58,59^. The shift towards more IgD-expressing (pre-GC) B cells and less IgA^+^ B cells can be explained by the reduced fraction of GC B cells observed in the cancerous colon of hypo-innervated mice. This scenario is supported by results obtained with *in vitro* cultured human GC B cells that responded vigorously to epinephrine stimulation in terms of cell number while sustaining a GC and surface IgA^+^ B cell phenotype coupled to a slight increase in the frequency of CD38^high^CD10^-^ plasma-cell like cells. An increase in the fraction of surface IgA^+^ B cells was counteracted by a corresponding reduction in secreted IgA, suggesting interference with full execution of an activated-B-cell-to-plasma cell differentiation program. Epinephrine stimulation had a remarkable positive influence on IgE class-switching at the expense of reduced IgG1, both isotypes being under the influence of Th2 cytokines^60^. To which extent this skewing exerts a negative impact on local protective IgG1-driven anti-CRC responses awaits future investigation. These results suggest that neurons play an important role in colon B cell homeostasis, function and terminal differentiation in contexts linked to chronic inflammation and tumorigenesis^61,62^.

While T cells have received strong attention to explain anti-cancer immunity, also B cells have recently been shown to actively modulate anti-tumor immune responses. Their role can be anti-tumorigenic, by modulating T cell priming, and producing antibodies targeting tumor cells, thereby promoting tumor cell elimination by NK cells^63^. However, B cells can also produce tumor-modulating factors (such as chemokines, cytokines and antibodies) and acquire regulatory functions capable of suppressing T cell cytotoxic activity, thereby sustaining tumorigenesis. IgG1 secretion has been shown to mediate tumor cytotoxicity, and was decreased in the neuronally-stimulated B cells, while IgE is linked to suppression of tumor cytotoxicity, and was increased after adrenergic stimulation^64^. IgA shows a more complicated profile as IgA-expressing B cells were increased, but IgA secretion was reduced by epinephrine treatment. This might indicate that the IgA-expressing B cells have not fully matured into IgA-secreting cells after four days of treatment, but are potentially able to boost IgA secretion on the long-term, which can have both pro- and anti-tumor effects^65^. Next to alterations in Ig secretion, IL-10 secretion from B cells increased after adrenergic stimulation. As this anti-inflammatory cytokine is known to be involved in B cell survival, class-switching and antibody production^66,67^, this result directly links with the increased proliferation and maturation of the epinephrine-treated B cells. The tumor immune surveillance role of IL-10 is mostly indirect and probably involves T cells in the TME^68^. Although frequency of different T cell subsets did not appear to be affected in hypo-innervated CRC mice, future experiments will be required to closely study their function and activity.

Cell network investigations based on transcriptome analyses point to distinct signaling pathways defining potential interactions between neurons, B cells and the tumor, involving extracellular matrix genes, EGFR-ligands, and lymphocyte regulators. Although these factors may have a direct effect on cancer cells, such as Granzyme B (Gzmb), which provides B cells with a cytotoxic capacity to directly kill tumor cells^69,70^, neuron-stimulated B cells may also influence the biology of other components of the TME via these factors. For example, B cells have been shown to induce collagen secretion from fibroblasts^71^, to a degree that is proportional to their plasma cell commitment state^72^, a state that we observed in *in vitro* cultured GC B cells upon epinephrine stimulation. Collagen was identified to promote both stemness and metastasis in CRC, thereby contributing to establish a pro-tumorigenic TME^73,74^. Moreover, the EGFR-ligand epiregulin (EREG), which was identified as communicator between neuronally-stimulated B cells and tumor cells, has been shown to promote tumor growth, with a common involvement of tumor-associated fibroblasts^75^. Interestingly, high *EREG* expression was associated with a better response to cetuximab treatment in metastatic CRC patients^76^. Other predicted factors, CXCL13 and CD7, have been identified to play a role in B cell recruitment as well as T cell immune infiltration of tumors and can thereby affect both tumor immunity and treatment response^77,78^. While CCL17 signaling from B cells can affect T cell infiltration, its pro-tumorigenic effect has also been linked to microbiota composition, which is a major contributor to the colonic TME^79,80^. Overall, the predicted interactions between epinephrine-treated B cells and CRC cells can affect multiple TME factors as well as tumor cells, with most effects linked to the establishment of a pro-tumorigenic environment. This was confirmed by comparing the epinephrine-stimulated B cell signature with human CRC B cell expression profiles, as this signature mostly resembles tumor plasma cell populations and associates with worse survival for CRC patients. Collectively, because adrenergic neurons induce B cell proliferation and maturation, although seemingly incomplete and associated with a pro-tumorigenic function, we hypothesize that they acquire a more regulatory phenotype^81^.

Together, our study reveals a previously unrecognized sympathetic neuron - B cell communication axis in CRC, providing evidence for direct neuro-modulation of B cell maturation and activation, with implications for chronic colon inflammation and CRC biology and a potential impact on disease progression and therapeutic responses. Until now, immune-related cancer studies have primarily focused on the presence of immune cell infiltration within tumors and the expression of immune checkpoint molecules. We propose that additional classification for patient prognosis and stratification would benefit from the inclusion of markers for B cells and neurons, and should consider their spatial relationship.

## Methods

### Mice

*Wnt1-Cre2* (022137; 129S4 background) and *Hand2^fl^*(027727; 129S1 background) transgenic mice were obtained from Jackson Laboratories. *Wnt1-Cre2* mice were crossed with *Hand2^fl/fl^* mice to generate *Hand2^fl/+^;Wnt1-Cre2* (hypo-innervated) and *Hand2^fl/+^*(control) littermates for experiments. All mice were characterized by genotyping PCR (primer sequences in Supplementary Table 1). Animals were housed with littermates in a maximum group size of five with *ad libitum* access to water and food. All animal experiments were conducted with approval from the Committee of Animal Welfare of Maastricht University and performed according to Dutch regulations (AVD1070020174386, AVD10700202215867).

### Immunofluorescence on myenteric plexus preparations

Colons from 12-week old mice (without cancer induction) were dissected as previously described^82^. Briefly, colons were isolated from cecum to rectum and cut open along the mesentery. Tissue was pinned flat in a Sylgard® coated Petri dish containing PBS and the mucosal and submucosal layers were peeled off using forceps, revealing the myenteric plexus. Tissue preparations were fixated in 4% PFA in PBS for 30 min at RT before immunofluorescent antibody staining. Myenteric plexus preparations were permeabilized with 1% Triton X-100 in PBS containing 4% donkey serum (Jackson ImmunoResearch; 017-000-121) for 2h. Primary antibody were incubated for 24h at 4°C and after washing, secondary antibodies were applied for 2h at 4°C (Supplementary Table 2). Hu-stained tissue preparations of proximal and distal sections of the colon were imaged on a Leica TCS SP8 confocal microscope (Fluotar VISIR; 25x, H_2_O immersion lens, NA = 0.95) and TH-stained tissue preparations of distal colon were imaged on a Zeiss LSM900 Airyscan 2 (Plan-ApoChromat; 20x, air immersion lens, NA = 0.55). At least two regions (tilescans of approximately 2.0 mm² total surface) per tissue preparation were imaged. Hu^+^ neurons were manually counted using the Cell Counter plugin in the ImageJ software (NIH, Bethesda, MD). TH immunoreactivity was quantified as proportion of signal to image area using maximum intensity projections of z-stack tilescans, and inversion and particle analysis plugins in the ImageJ software. Cell number or area percentage per region was averaged per preparation and normalized for colon stretching during dissection.

### Experimental model of colorectal cancer and inflammation

To induce colitis-associated CRC, *Hand2^fl/+^* and *Hand2^fl/+^;Wnt1-Cre2* (10-12 weeks old males (unless otherwise specified)) were injected intraperitoneally with a single dose of 10 mg/kg of the carcinogen azoxymethane (AOM; sc-358746A, Santa-Cruz) dissolved in sterile phosphate-buffered saline (PBS). Dextran sodium sulfate (DSS; 36,000–50,000 M.Wt., Colitis Grade, MP Biomedicals) was dissolved in autoclaved tap water (1.5%-2%, w/v) and supplied in the drinking water for five days during the second, fifth (and eighth) week of the protocol. Mice had access to autoclaved tap water for the remainder of the protocol. Mouse weights were monitored weekly over the duration of the experiment. After 10-12 weeks, mice were euthanized using CO_2_ asphyxiation, and their organs were collected for further analysis. To induce colitis, *Hand2^fl/+^*and *Hand2^fl/+^;Wnt1-Cre2* (10-12 weeks old males) were given DSS (3%) in the drinking water for seven days and subsequently euthanized using CO_2_ asphyxiation, collecting their organs for further analysis.

### *In vivo* tumor tracking

Computed tomography (CT) scans were acquired in week 3, week 5, week 7, and week 9 of the CRC induction protocol. Mice were fasted for 16h before the procedure and anesthetized using isoflurane (IsoFlo®) during the procedure. Contrast enhancement was adapted from a previously described method^83^. Briefly, 1.5 ml iomeprol (Iomeron® 350; diluted 1:5 in sterile PBS) was injected intraperitoneally and 1.5 ml sodium ioxitalamate (Telebrix® Gastro, Guerbet; diluted 1:10 in sterile PBS) rectally. CT scans were acquired on an X-RAD 225Cx^84^ small animal irradiator (Precision X-ray, Inc)^85^ at 80 kVp, 3 mA, 120s, 39cGy/scan under isoflurane anesthesia. The Feldkamp^86^ filtered back projection (Pilot; Precision X-ray, Inc) method was used to reconstruct image projections. Visualization and image analysis were performed using SmART-ATP software (v. 2.1, SmART Scientific Solutions BV, Maastricht, The Netherlands). Tumor volumes were calculated by contouring the tumor manually in every 2D plane and calculating the 3D tumor volume as previously described^87^.

### Sample preparation for staining and RNA sequencing

After sacrifice, colons were cut open along the mesentery to count tumor number and measure tumor size. One tumor with an approximate size of 2 mm was collected per mouse, snap frozen in liquid nitrogen, and stored at −80°C for subsequent RNA isolation. The remaining tissue was Swiss-rolled^88^, stored in a tissue processing cassette, and fixed in 4% paraformaldehyde (PFA) for at least 36h at RT. Swiss rolls were then sequentially immersed in 15% and 30% (wt/vol) sucrose at 4°C overnight or until sunk for cryoprotection. Tissue-Tek OCT (Sakura) was used to embed the Swiss rolls and 7 µm cryosections were cut on a cryostat (Leica CM3050 S).

### Immunohistochemistry and immunofluorescence on Swiss rolls

Cryosections of colon Swiss rolls were stained with hematoxylin and eosin (H&E), or immunohistochemically stained (Supplementary Table 2). Cryosections were first incubated with 0.3% hydrogen peroxide in methanol for 20 min to quench endogenous peroxidase activity. Nonspecific antibody binding was blocked by incubation with 20% fetal calf serum in PBS Tween 0.1%. Sections were incubated with primary antibody diluted in PBS, 1% BSA, 0.1% Tween (pH 7.2-7.6) for 1h at RT. Following incubation with secondary IgGs antibody, the bound antibodies were visualized with 3,3’-diaminobenzidine (DAB, Dako) or Vector Red (VectorLabs) substrate. For double staining, the antibody incubations were repeated with a different primary and secondary antibody and visualization substrate. Slides were counterstained with hematoxylin, dehydrated, and mounted. Images were acquired using a Leica DM300 LED microscope equipped with a Leica DFC320 camera (Leica Microsystems) and the QWIN V3 software (Leica) or scanned with the 3DHISTECH Pannoramic1000 and visualized with CaseViewer version 2.4.0.119028.

For IF staining, cryosections of colon Swiss rolls were permeabilized with 1% Triton X-100 in PBS containing 4% donkey serum (Jackson ImmunoResearch; 017-000-121) for 2h. Primary antibody incubation was performed overnight at 4°C and after washing, secondary antibodies were applied for 2h at 4°C (Supplementary Table 2). Colon Swiss rolls were imaged on a Leica DM5000B microscope (10× magnification).

### Tumor classification

Tumors were identified on H&E and beta-catenin-stained Swiss-roll sections by a trained animal pathologist, and classified based on histology and following Boivin et al.^89^. Tumors were categorized as hyperplasia, low grade adenoma, high grade adenoma, or adenocarcinoma. In brief, (I) hyperplasia was characterized by gross thickening of the mucosa; (II) low grade adenoma by a circumscribed neoplasm, branching or elongation of crypts, and presence of mucous secretion; (III) high grade adenoma by a marked reduction of interglandular stroma and a complex irregularity of glands, numerous mitoses with abnormal mitotic figures, and absence of mucus secretion; and (IV) adenocarcinoma as malignant neoplasia that is infiltrating in blood vessels or penetrating the muscularis mucosae.

### Transmission electron microscopy

Human normal colon and tumor tissue samples from CRC patients were cut in fine pieces and prepared for transmission electron microscopy. Samples were fixed with Karnovsky’s fixative (2.5% glutaraldehyde and 2% paraformaldehyde in 0.1M sodium cacodylate; pH7.4), washed with 0.1M sodium cacodylate buffer and post fixed with 1% osmium tetroxide in 0.1M sodium cacodylate buffer containing 1.5% potassium ferricyanide for 1h in the dark at 4°C. Samples were dehydrated in ethanol, infiltrated with Epon resin for two days, embedded in the resin, and polymerized at 60°C for 48h. Ultrathin sections of 70 nm were obtained using a Leica Ultracut UCT ultramicrotome (Leica Microsystems) and mounted on Formvar-coated copper grids. Tissues were stained with 2% uranyl acetate in 50% ethanol and lead citrate. Sections were imaged with a Tecnai T12 Electron Microscope equipped with an Eagle 4kx4k CCD camera (Thermo Fisher Scientific).

### Tumor RNA sequencing and data analysis

Tumor RNA was isolated from *Hand2^fl/+^* and *Hand2^fl/+^;Wnt1-Cre2* mice using the RNeasy Mini kit (Qiagen) according to manufacturer’s instruction. In brief, tissue was lysed in 350 µl RLT buffer containing 1% β-mercaptoethanol and genomic DNA was removed by incubating with DNase I solution. Isolated RNA was dissolved in 60 µl RNase-free water and stored at −80°C. RNA sequencing analysis was performed at GenomeScan B.V. (Leiden, the Netherlands) on a NovaSeq6000. Samples were processed using the NEBNext Ultra II Directional RNA Library Prep Kit for Illumina® (#E7760S/L, New England Biolabs) according to protocol. Briefly, oligo-dT magnetic beads were used to isolate mRNA from total RNA, and mRNA was subsequently fragmentized and used for cDNA synthesis. This was used for ligation with sequencing adapters and PCR amplification. Sample quality and concentration were determined on a Fragment Analyzer and all samples passed quality control before sequencing. Data analysis was performed by ServiceXS™ GenomeScan. Image analysis, base calling, and quality check were performed with the Illumina data analysis pipeline RTA3.4.4 and Bcl2fastq v2.20. Gene set enrichment analysis (GSEA) was performed using the msigdbr 7.5.1 R package. The input for this analysis was an ordered list of genes and the set of H pathways from MsigDB. The fgsea function in the msigdbr package was applied to the gene list and pathway set, and the resulting enrichment scores and corresponding *P* values and BH-adjusted *P* values were calculated. Another GSEA was performed for hallmarks of cancer^44^. Hallmarks of cancer gene lists were retrieved from https://figshare.com/articles/dataset/Hallmarks_of_Cancer_Gene_Set_Annotation/4794025?file=7881835. The human hallmarks of cancer gene lists were then converted to mouse orthologs using the orthologs dataset from Jax Labs. Gene set enrichment analysis (GSEA) was conducted on the differentially expressed genes for each set of hallmarks of cancer genes using Fisher’s exact test (fisher.test R function, stats package version 4.2.0), as previously described^32^. False discovery rate (FDR) *P* value adjustment was applied to correct for multiple testing. Volcano plots were generated using the EnhancedVolcano 1.14.0 R package.

### Immune cell isolation and flow cytometry

Cells were isolated from the colon of healthy, DSS-treated and AOM/DSS-treated *Hand2^fl/+^* and *Hand2^fl/+^;Wnt1-Cre2* mice as described previously^38^. In brief, mesenteric fat was removed, the colon was dissected open longitudinally, feces were removed and number and size of tumors was assessed. Tumors were removed from the colon and the colon was cut into 1 cm pieces. Tissues were then stored in cRPMI, containing 10% fetal calf serum (FCS), 1 mM sodium pyruvate (Gibco), 1× non-essential amino acids (Gibco), and PSEPx (1% pen/strep (Lonza), 25 µg/ml enrofloxacin (Thermo Scientific), 100 units/ml polymyxin B (Gibco) in RPMI-1640, on ice before further processing. Colons were washed at 200 RPM at 37°C for 20 min in 5 mM DTT containing wash buffer (2% FCS and PSEPx-containing Hank’s buffered salt solution; HBSS) to remove mucus and subsequently washed three times 15 min in 5 mM EDTA-containing wash buffer to remove epithelial cells. Colon tissues were minced into fine pieces and digested in 0.2 Wünsch units/ml Liberase TM (Roche) and 5 mg/ml DNAse I (Roche) in HBSS for 30 min at 37°C while shaking at 200 RPM. The enzymatic reaction was stopped by storing the samples on ice and adding 0.5 volume cRPMI. Samples were homogenized by triturating through an 18-gauge needle and filtered through a 100 µm cell strainer (Greiner). All cells were washed in 2% FCS in HBSS and pelleted for subsequent staining. Antibodies used for flow cytometry staining are listed in Supplementary Table 2. Data were acquired on a 4 laser (UV/V/B/R) Cytek Aurora spectral cytometer using SpectroFlo® analysis software. Fluorescence minus one controls were used to determine gating strategy.

### Microbiome analysis

Stool was collected from *Hand2^fl/+^* and *Hand2^fl/+^;Wnt1-Cre2* animals before cancer induction (10-12 weeks of age) and after cancer induction, the day before euthanasia. To collect stools, the animals were put in a sterile 1L beaker until defecation. Stools were collected sterile, snap frozen in liquid nitrogen, and stored at −80°C until analysis. Microbial DNA was isolated from one or multiple stools per mouse using the QIA Fast DNA Stool Mini Kit (Qiagen) according to manufacturer’s instructions. For the analysis of bacterial metagenomes on species level (all executed by GenomeScan B.V.), the V4 region of the microbial 16S ribosomal RNA (rRNA) was analyzed using a nested PCR including dual index Illumina compatible primers. Next generation sequencing of the 16S rRNA amplicon (Illumina NovaSeq 6000), quality control (FastQC) and trimming of reads (Cutadapt) were performed, followed by inferring Amplicon Sequence Variants (ASVs) using DADA2, and taxonomical classification by QIIME2 workflow. An average of 532,235 (375,478–700,489) reads per sample was obtained. Unwanted taxa (mitochondria and chloroplast) were excluded, rarefaction curves and alpha and beta diversity indices were generated, and differentially abundant taxa (ANCOM) were identified.

### SCENITH

Single cell metabolism (SCENITH) protocol was performed on tumors from AOM/DSS-treated *Hand2^fl/+^* and *Hand2^fl/+^;Wnt1-Cre2* mice^90^. Cell isolation of the flow cytometry protocol was largely followed. However, all steps and storage times were done at RT. After transfer of the single cell suspension to V-shaped 96-well plates (five wells per sample), plates were incubated for 30 min at 37°C and 5% CO_2_ to ensure normal metabolism. Once incubated, 5 µL of inhibitors were added to the wells divided in five conditions: Control (Co), Deoxy-D-Glucose (DG, final concentration 100mM), Oligomycin A (O, final concentration 1μM), DG+O, and Harringtonine (H, final concentration 2 μg/ml). To all wells, 5.3 µL of prediluted (20x) Puromycin (final concentration 10 μg/ml) was added, after which the samples were mixed and incubated at 37°C and 5% CO_2_ for 40 min. Subsequently, cells were washed, stained to identify tumor cells and samples were acquired on a 4 laser (UV/V/B/R) Cytek Aurora spectral cytometer using SpectroFlo® analysis software (Supplementary Table 2 and Supplementary Figure 3A). To determine metabolic dependencies and capacities of the tumor cells, formulas with median fluorescence intensity values of the puromycin signal for the different conditions were used (Supplementary Figure 3C).

### Primary human B cell assays

Germinal center B cell enriched human primary cultures of tonsil tissues were obtained as previously described^39^. B cells were cultured on YK6-CD40Lg-IL21 feeder cells in Advanced RPMI 1640 (Gibco) supplemented with 20% FCS, 1% Pen/Strep and 1% L-glutamate for a maximum of 10 days with passaging on new feeder cells every 3-4 days. Cultures were treated with the neurotransmitter acetylcholine (200 μM; Sigma-Aldrich), epinephrine (100 μM; Sigma-Aldrich) or VIP (500 nM; Sigma-Aldrich). At multiple timepoints, cells were harvested for cell counting, cell viability assay, and flow cytometry. Cell counting was performed on a Countess Automated Cell Counter (Invitrogen) using tryphan blue to differentiate between live and dead cells. Cell viability was examined by CellTiter-Glo luminescent assay (Promega) as by manufacturer’s instructions and read-out using EnVision 2104 multilabel reader (PerkinElmer). Flow cytometry was executed by staining the B cells with a B cell subtype marker panel and acquired on a 5 laser (UV/V/B/YG/R) Cytek Aurora spectral cytometer using SpectroFlo® analysis software (Supplementary Table 2 and Supplementary Figure 6A).

### Secretome analysis

Conditioned medium from untreated and neurotransmitter-treated B cells was harvested and stored at −80°C. LEGENDplex secretome analysis was performed using the human immunoglobulin isotyping panel (8-plex) and human B cell panel (13-plex) (Biolegend) and following manufacturer’s instructions. Shortly, 20-25 μL conditioned medium was incubated consecutively with assay beads for 2h, biotinylated detection antibody for 1h, and Streptavidin-PE for 30 min. Data were acquired by BD FACSCanto II using FlowJo analysis software.

### Bulk B cell RNA sequencing and data analysis

Total RNA was extracted from B cell pellets using Maxwell® RSC miRNA Tissue Kit (Promega). RNA quantification was performed by Nanodrop 1000 (Agilent) and Qubit 3.0 Fluorometer (Life Technologies) and RNA integrity was evaluated with Bioanalyzer 2100 (Agilent) using the RNA 6000 Nano Kit (Agilent). Libraries were prepared with the Illumina TruSeq RNA Sample Prep Kit and sequenced on the NextSeq2000 to get single-end 150 bp reads. Conversion to fastq was performed using the Dragen bcl-convert pipeline. The reads were aligned to hg38 using the splice-aware MapSplice tool and the resulting BAM files underwent processing via ubu sam-xlate and sam-filter to translate genomic into transcriptomic coordinates and to exclude reads with indels, large inserts and zero mapping quality. The quantification of transcripts and genes was conducted using RSEM and GENCODE v33. PCA (Principal Component Analysis) was performed to identify the specific transcriptomic profiles of each treatment group. Differently expressed genes (DEG) were calculated by “DEseq2” R package and adjusted P-value<0.05 and log2 Fold Change>1 or <-1 were used as threshold for high and low expressed genes. Gene Ontology (GO) analysis was performed using the high expressed genes with the R package “ClusterProfile” with the threshold adjusted P-value<0.05.

### Integration of single-cell RNA sequencing data

Single-cell RNA sequencing datasets were obtained from GEO (GSE132465) and ArrayExpress (E-MTAB-8107, E-MTAB-6149, E-MTAB-6653). Raw gene expression matrices and metadata were processed using the Seurat R package (v4.2.2)^91^. Low-quality cells were excluded based on the following thresholds: >200 and <4000 detected genes, <30,000 total RNA molecules, and <25% mitochondrial gene content. After filtering, 16,404 cells from non-tumor and 47,285 cells from tumor samples were retained. Quality-controlled data were normalized using 3,000 highly variable genes, followed by dimensionality reduction with the top 20 principal components (PCs). Harmony was applied to correct for batch effects across patients^92^. UMAP was used for nonlinear dimensionality reduction, and clustering was performed using Seurat’s graph-based approach (FindNeighbors, FindClusters; resolution=0.5). To enable consistent comparisons, 1,000 cells per cluster were randomly downsampled. Differential gene expression (DGE) analysis was conducted using the Wilcoxon rank-sum test (adjusted P<0.05), and initial cell type annotation relied on canonical marker genes. B and plasma cells were subsequently isolated for subpopulation analysis. UMAP (20 dimensions) and clustering (resolution=0.3) revealed three B cell and four plasma cell clusters. For comparison with bulk RNA-seq, samples were converted to Seurat objects and integrated using canonical correlation analysis (CCA) anchoring. Cluster-specific marker genes (log₂ fold-change>0.25, adjusted P<0.05) were selected for further analysis.

Transcriptomic similarity between bulk and single-cell populations was assessed using Spearman correlation of normalized expression values, and results were visualized as hierarchically clustered heatmaps. For ligand prediction, NicheNet was used with genes upregulated in tumor epithelial cells (vs. non-tumor) as targets^93^. Ligands enriched in epinephrine-treated B cells (vs. untreated) from bulk RNA-seq were filtered and matched to the NicheNet output.

### TCGA RNA-seq data analysis

Expression data and clinical information data were downloaded from TCGA-COAD by “TCGAbiolinks” R packages. After matching the clinical information and expression data, high expression genes in epinephrine-treated B cells compared to untreated B cells were used as the Epi risk assessment gene list, and PCA was applied on each patient with their Epi risk gene expression. To identify the optimal cut-off value for stratifying patients into an Epi low and high group based on their continuous Epi risk score, the surv_cutpoint() function from the “survminer” R package was applied by computing maximally selected rank statistics (max log-rank test statistic) across all possible cut-off points. The analysis was performed with the parameters minprop=0.20 to ensure that each resulting group contains at least 20% of the total samples, thereby avoiding extremely unbalanced stratification. The optimal cut-off value was then used to divide patients into “high-Epi risk” and “low-Epi risk” groups. Differences in clinicopathological characteristics between subgroups based on the Epi risk score were evaluated using t-tests or chi-square tests where appropriate. Overall survival (OS) was defined as time from cancer diagnosis until death by any cause, or end of follow-up and was restricted at 6 years of follow-up. Univariable survival analyses to assess the association between the Epi risk score and overall survival (OS) were performed using Kaplan-Meier and log-rank tests. Hazard ratios (HR) and corresponding 95% cinfidence intervals (CI) were determined with Cox proportional hazards models adjusted for the potential confounders sex and age. The proportional hazards assumption was tested using the scaled Schoenfeld residuals. All survival analyses were performed using STATA 17.0.

### Statistical analysis

Data were analyzed by two-tailed unpaired t-test, Mann-Whitney *U* test (for non-parametric data) or one-way ANOVA with post-test correction (Bonferroni or Dunnett; for multiple comparisons) using SPSS version 25 and/or GraphPad Prism5 (version 5.04). P-values and stars represent the significance level: * P<0.05, ** P<0.01, *** P<0.001, **** P<0.0001.

## Acknowledgements

We thank Dr. Tullia Bruno for her expertise and suggestions to improve the manuscript. We acknowledge the Microscopy CORE lab at Maastricht University and the AOMC at Hasselt University (funded by The Research Foundation Flanders (FWO) [I001222N]) for support with microscopy experiments. This study was supported by a BOF-mandate from Hasselt University (BSFWBIOMED) and Maastricht University Medical Centre+ (M.S.T.), Kankeronderzoeksfonds Limburg (KOFL) (M.S.T.), a long-term travel grant by FWO [V472323N (M.S.T.)], and the Dutch Research Council (NWO) VENI grant [016.186.124 (V.M.)] and VIDI grant [09150172110100 (V.M.)].

## Author Contributions

V.M. and W.B. designed and supervised the study. M.S.T. and S.L.S. designed and supervised functional experiments. M.S.T., S.L.S., L.C., J.dV., J.V., N.V., S.S., K.W., S.S. performed experiments and collected data for this study. M.S.T., S.L.S., Y.Z., M.I., J.dV., K.W., K.S. performed data analyses. M.J.G. was involved as experimental animal pathologist. L.T. and E.W. were involved in flow cytometry experiments. F.V. and L.D. provided software and expertise for the in vivo CT scanning experiments and analyses. C.L.I. and W.vdW. advised on and performed the electron microscopy. S.C. and A.B. provided the in vitro B cell culture models and experimental framework. M.S.T., S.L.S., W.B. and V.M. drafted the manuscript. All authors have assisted with result interpretation and revised the manuscript.

