## Supplementary Tables and Figures for "Adrenergic signaling induces a pro-tumorigenic B cell state in colorectal cancer"

**Supplementary Table 1| PCR Primers for genotyping**

| **Target** | **Forward primer** | **Reverse primer** | **Annealing temp.** |
|---|---|---|---|
| Hand2$^{fl/fl}$ | ACTTGCTGACTGGGTCCTTG | CTCGGCCTAGAGGACACTGA | 60°C |
| Wnt1-Cre2 | CAGCGCCGCAACTATAAGAG | CATCGACCGGTAATGCAG | 60°C |
| Internal pos. ctrl | CAAATGTTGCTTGTCTGGTG | GTCAGTCGAGTGCACAGTTT | 60°C |

**Supplementary Table 2| List of antibodies used for immunohistochemistry, immunofluorescence, and flow cytometry**

| Antibody target | Conjugate | Host | Dilution | Supplier | Cat. no. |
|---|---|---|---|---|---|
| **Immunohistochemistry** | | | | | |
| B catenin | - | Rabbit | 1:75 | Cell Signaling Technologies | 8480 |
| B220 | - | Rat | 1:500 | R&D | MAB1217 |
| Tubulin b-3 | - | Rabbit | 1:500 | Biolegend | 802001 |
| Rat IgG | HRP | Goat | 1:200 | Invitrogen | 31470 |
| Rabbit IgG | AP | Goat | Ready-to-use | Immunologic | DPVB55AP |
| Rabbit IgG | HRP | Goat | Ready-to-use | ImmunoLogic | DPVR110HRP |
| **Immunofluorescence** | | | | | |
| HuC/D | - | Mouse | 1:200 | Thermofisher | A-21271 |
| TH | - | Rabbit | 1:400 | Abcam | Ab112 |
| B220 | - | Rat | 1:500 | R&D | MAB1217 |
| Tubulin b-3 | - | Rabbit | 1:500 | Biolegend | 802001 |
| Rat IgG | Alexa 594 | Goat | 1:1000 | Invitrogen | A21471 |
| Rabbit IgG | Alexa 488 | Donkey | 1:1000 | Thermofisher | A21206 |
| Mouse IgG | Alexa 594 | Donkey | 1:1000 | Thermofisher | A-21203 |
| **Flow Cytometry** | | | | | |
| LIVE/DEAD™ | Aqua | Rat | 1:1000 | Invitrogen | 10321873 |
| FcBlock (CD16/CD32) | - | Rat | 1:100 | Invitrogen | 14-0161-85 |
| CD45 | PerCp | Rat | 1:100 | Biolegend | 103130 |
| CD3 | V450 | Rat | 1:100 | Invitrogen | 48-0032-82 |
| CD3 | BUV496 | Hamster | 1:50 | BD Bioscience | 612955 |
| CD4 | APC-H7 | Rat | 1:100 | BD Bioscience | 560181 |
| CD8 | FITC | Rat | 1:50 | Invitrogen | 11-0081-85 |
| CD45R (B220) | SB702 | Rat | 1:100 | Invitrogen | 67-0452-82 |
| CD45R (B220) | APC | Rat | 1:50 | Invitrogen | 17-0452-82 |
| CD25 | APC | Rat | 1:100 | Invitrogen | 17-0251-81 |
| CD25 | PE-Cy7 | Rat | 1:300 | Invitrogen | 25-0251-82 |
| FoxP3 | PE | Rat | 1:40 | Invitrogen | 12-5773-82 |
| MHCII | FITC | Rat | 1:100 | Invitrogen | 11-5322-82 |
| CD11b | eFluor450 | Rat | 1:300 | Invitrogen | 48-0112-80 |
| CD11b | PE | Rat | 1:800 | Invitrogen | 12-0112-82 |
| LY6C | APC | Rat | 1:100 | Miltenyi | 130-123-769 |
| LY6G | APC-Cy7 | Rat | 1:100 | BD Bioscience | 560600 |
| F4/80 | BV421 | Rat | 1:100 | Biolegend | 123137 |
| CD170 (Siglec-F) | PE | Rat | 1:300 | Invitrogen | 12-1702-80 |
| CD64 | PerCP-eF710 | Rat | 1:100 | Invitrogen | 46-0641-80 |
| CD11c | PE-Cy7 | Rat | 1:100 | Invitrogen | 25-0114-82 |
| CD19 | BV421 | Rat | 1:100 | Biolegend | 115549 |
| CD19 | PE-Cy7 | Mouse | 1:100 | eBioscience | 25-0199-42 |
| IgA | PE | Rat | 1:50 | Invitrogen | 12-4202-81 |
| IgA | AlexaFluor647 | Goat | 1:20 | Jackson | 109-606-011 |
| IgM | APC | Rat | 1:100 | BD Bioscience | 550676 |
| IgD | eFluor450 | Rat | 1:200 | Invitrogen | 48-5993-82 |
| IgD | FITC | Goat | 1:160 | Invitrogen | H15501 |
| IgG | PerCP | Goat | 1:40 | Jackson | 109-126-098 |
| GL7 | Pe-Cy7 | Rat | 1:50 | Biolegend | 144619 |
| CD95 | PerCP-eF710 | Mouse | 1:50 | Invitrogen | 46-0951-82 |
| CD43 | FITC | Rat | 1:300 | Biolegend | 143203 |
| CD5 | APC-eF780 | Rat | 1:300 | Invitrogen | 47-0051-80 |
| CD80 | BV605 | Hamster | 1:200 | Biolegend | 104729 |
| PD-L1 | SB780 | Rat | 1:200 | Invitrogen | 78-5982-82 |
| Sca | PerCP-Cy5.5 | Rat | 1:500 | Invitrogen | 45-5981-80 |
| CD27 | PE | Mouse | 1:100 | Biolegend | 356406 |
| CD38 | PerCP-eF710 | Mouse | 1:40 | eBioscience | 46-0389-42 |
| CD10 | BV421 | Mouse | 1:10 | BD Bioscience | 562902 |
| EpCAM | FITC | Rat | 1:20 | Invitrogen | 11-5791-82 |

**Supplementary Table 3| Clinicopathological characteristics of the included TCGA COAD patients in the total cohort as well as divided for low and high Epinephrine B cell signature.**

| Patient demographics | Total (N=391) | Epi low (N=252) | Epi high (N=139) | P-value |
|---|---|---|---|---|
| **Age at diagnosis, Mean ± SD** | 66.7 ±12.9 | 67.1 ± 12.3 | 65.9 ± 13.0 | 0.390 |
| **Sex, n (%)** | | | | |
| Male | 210 (54) | 139 (55) | 71 (51) | |
| Female | 181 (46) | 113 (45) | 68 (49) | 0.439 |
| **Cancer stage, n (%)** | | | | |
| Stage I | 68 (19) | 45 (19) | 23 (17) | |
| Stage II | 141 (39) | 91 (40) | 50 (38) | |
| Stage III | 102 (28) | 58 (25) | 44 (33) | |
| Stage IV | 52 (14) | 36 (16) | 16 (12) | 0.400 |
| **Histological type, n (%)** | | | | |
| Adenocarcinoma | 334 (87) | 219 (89) | 115 (83) | |
| Mucinous adenocarcinoma | 52 (13) | 28 (11) | 24 (17) | 0.101 |
| **History colon polyps, n (%)** | | | | |
| Yes | 118 (35) | 81 (37) | 37 (32) | |
| No | 217 (65) | 140 (63) | 77 (68) | 0.446 |
| **Vascular invasion, n (%)** | | | | |
| Yes | 152 (39) | 88 (35) | 64 (46) | |
| No | 239 (61) | 164 (65) | 75 (54) | 0.031 |
| **Lymph node positive ratio, Mean ± SD** | 0.10 ± 0.20 | 0.09 ± 0.17 | 0.13 ± 0.24 | 0.102 |

### Supplementary Figures

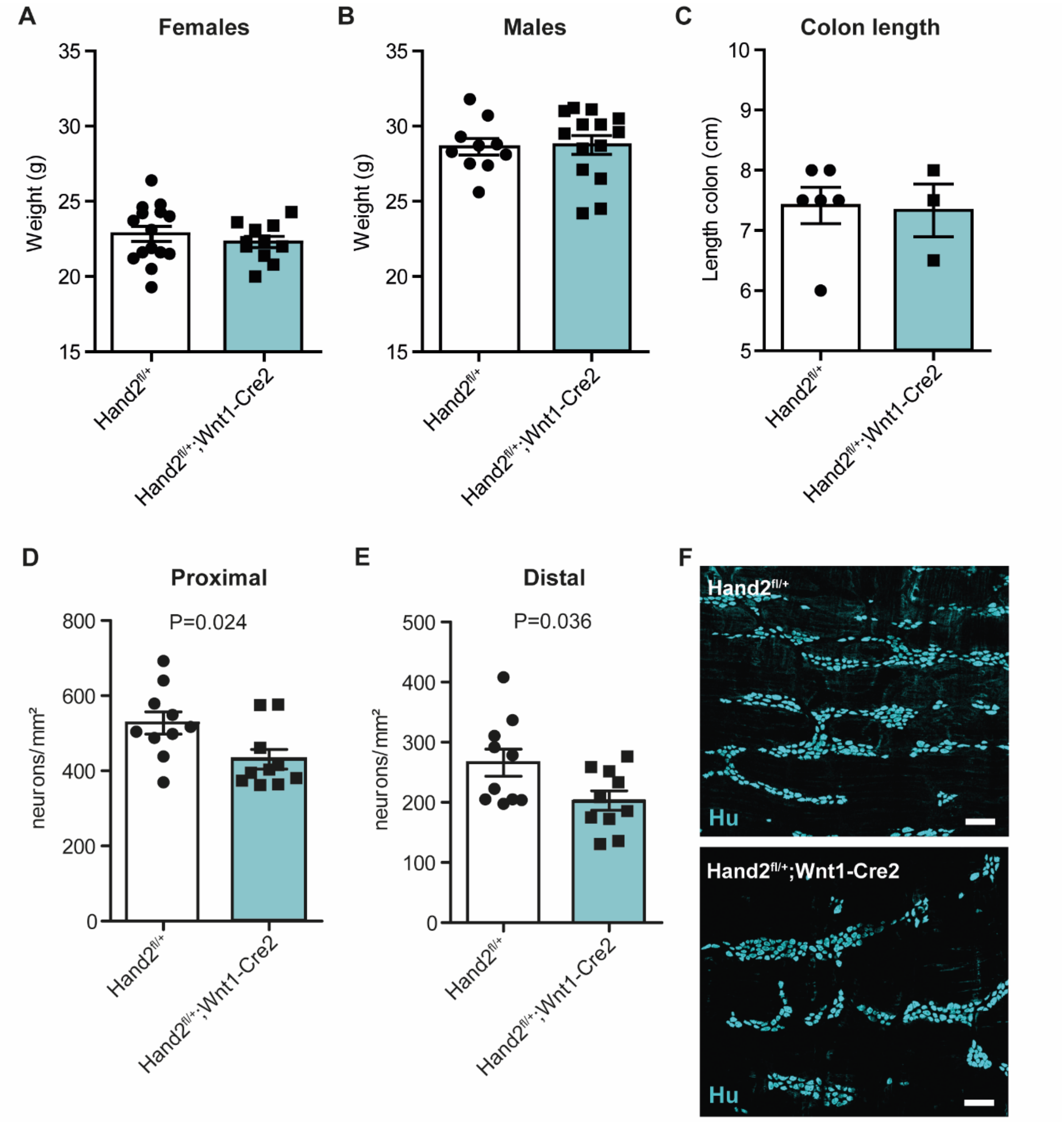


**Supplementary Figure 1| Healthy 11-week-old mice were characterized for body weight, colon length, and enteric neuron number.** Body weight did not significantly differ between **A|** female (N=15 and N=10) nor **B|** male (N=10 and N=14) *Hand2*$^{fl/+}$ and *Hand2*$^{fl/+}$*;Wnt1-Cre2* mice, nor did **C|** colon length differ (N=6 and N=3). The number of myenteric neurons was significantly reduced in *Hand2*$^{fl/+}$*;Wnt1-Cre2* (N=10) compared to *Hand2*$^{fl/+}$ (N=10) in both **D|** proximal and **E|** distal colon, as quantified by **F|** immunofluorescent Hu labeling. Scale bars are 100 µm.

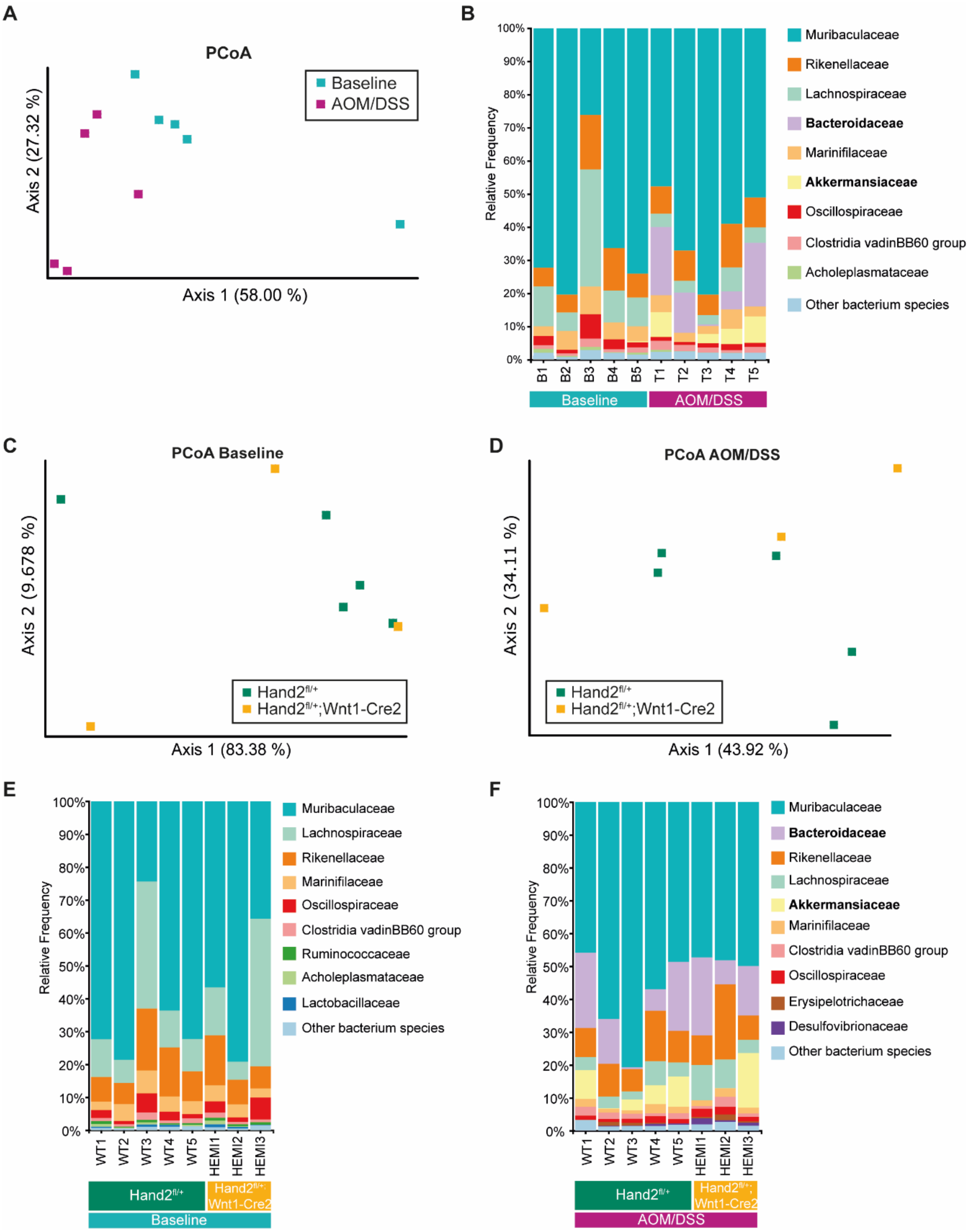


**Supplementary Figure 2| Fecal microbiome was assessed using 16S rRNA sequencing before (baseline) and after cancer induction (AOM/DSS). A|** Principal Component Analysis (PCoA) shows a significant (Weighted UniFrac *P*=0.008) difference in fecal microbiome composition between control mice at baseline and after AOM/DSS treatment. **B|** The bacterial families *Bacteroidaceae* and *Akkermansiacea* are significantly upregulated after AOM/DSS treatment. **C|** Principal Component Analysis (PCoA) shows no significant difference in the microbiome when comparing *Hand2*$^{fl/+}$ (N=5) and *Hand2*$^{fl/+}$*;Wnt1-Cre2* mice (N=3) at baseline and **D|** after AOM/DSS treatment. **E|** Relative microbiota frequencies are similar across *Hand2*$^{fl/+}$ and *Hand2*$^{fl/+}$*;Wnt1-Cre2* mice at baseline and **F|** after AOM/DSS treatment.

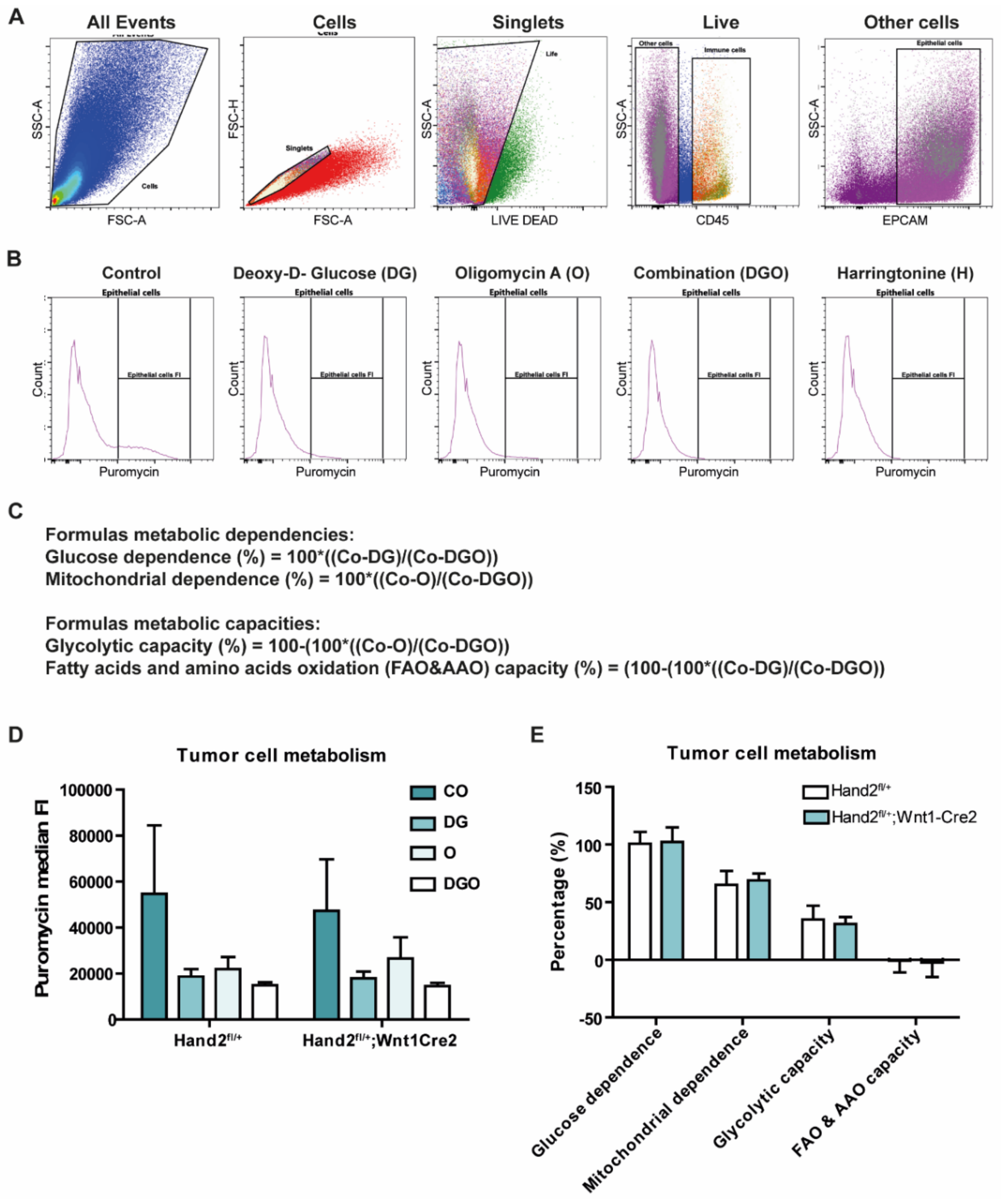


**Supplementary Figure 3| Single cell metabolism (SCENITH) set-up and analysis of tumors from *Hand2$^{fl/+}$* and *Hand2$^{fl/+}$;Wnt1-Cre2* mice. A|** Single cell metabolism analysis was performed on a Cytek® Aurora multispectral system, using a tumor cell panel. **B|** Puromycin signal, as a measure for protein synthesis and metabolic activity, decreases from control tumor epithelial cells to cells treated with deoxy-D-glucose (glycolysis inhibitor), Oligomycin (oxidative phosphorylation inhibitor) or a combination of both. Herringtonine (translation inhibitor) is used as negative control. **C|** Specific formulas were used to convert the puromycin signals from the different conditions to values for the glucose and mitochondrial dependences, and the glycolytic and fatty acids/amino acids oxidation capacities. **D|** Median Fluorescent Intensity (FI) of puromycin (as a measure for active metabolism) decreases most when glycolysis (DG) or both glycolysis and oxidative phosphorylation (DGO) are inhibited compared to control tumor cells (CO). **E|** The tumor cells are most dependent on glycolysis as metabolic pathway, but both glycolytic as well as mitochondrial dependence and capacity are not affected by hypo-innervation.

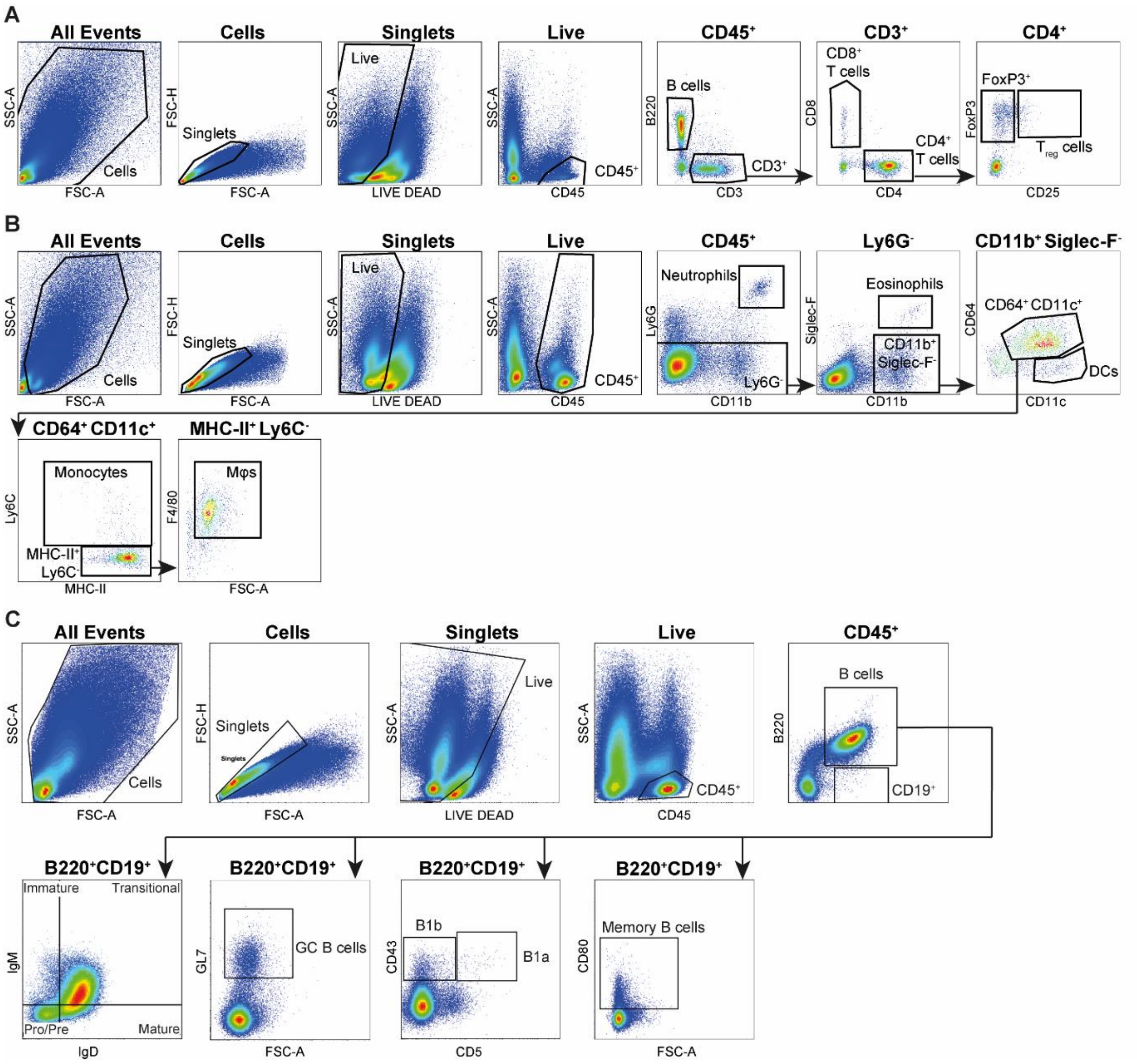


**Supplementary Figure 4| Gating strategy for flow cytometry analysis of healthy, inflamed and cancerous colon of *Hand2$^{fl/+}$* and *Hand2$^{fl/+}$;Wnt1-Cre2* mice.** Flow cytometry analysis was performed using a Cytek® Aurora multispectral system, with **A|** a lymphoid antibody panel, **B|** a myeloid antibody panel, and **C|** a B cell subtype panel.

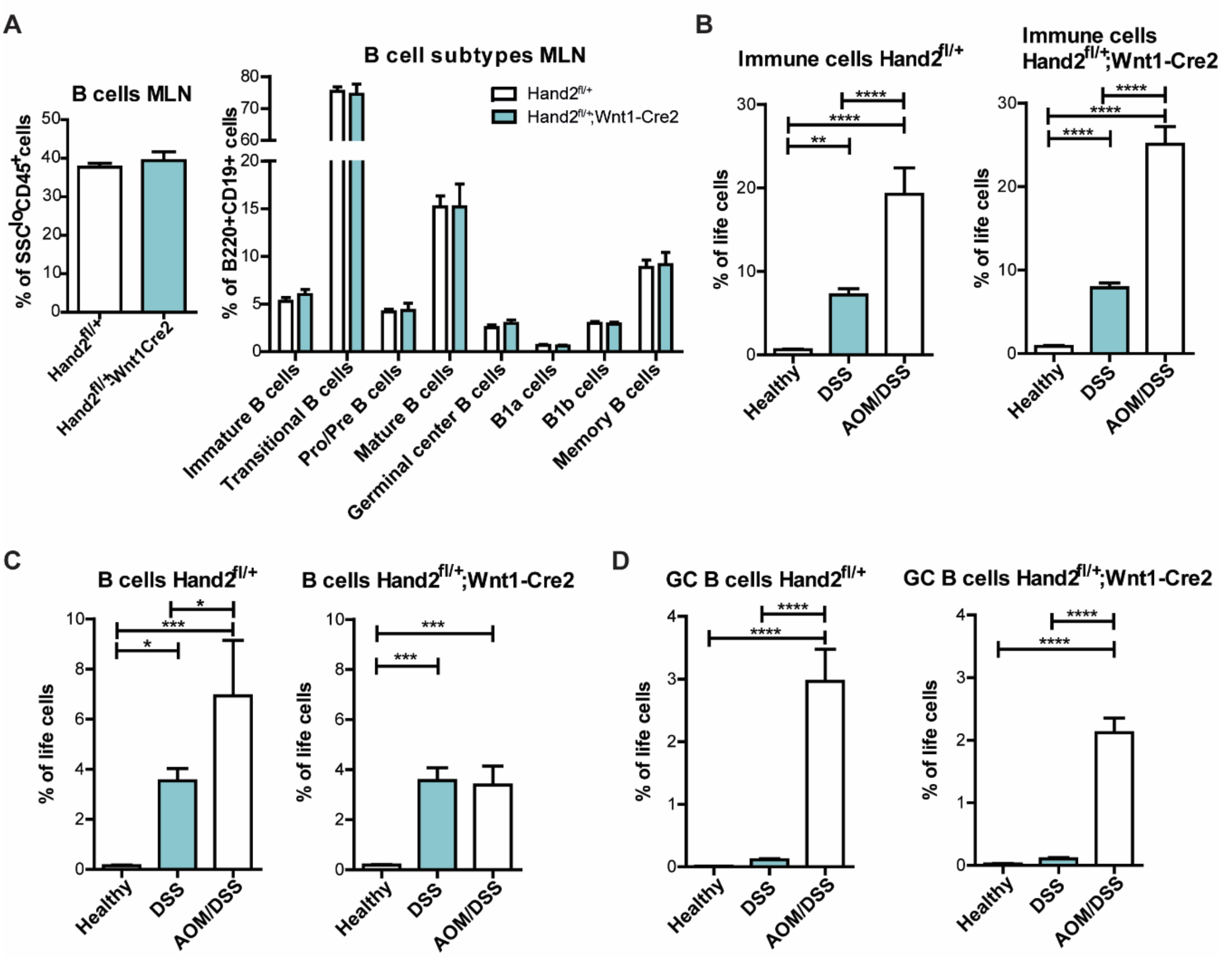


**Supplementary Figure 5| Flow cytometry analysis of mesenteric lymph nodes and comparisons between healthy, inflamed and cancerous colon. A|** B cells and B cell subtypes are not affected in the mesenteric lymph nodes (MLN) of *Hand2$^{fl/+}$;Wnt1-Cre2* (N=6) compared to *Hand2$^{fl/+}$* mice (N=6) as examined by flow cytometry. **B|** Immune cells increase significantly from healthy to inflamed to cancerous colon in both *Hand2$^{fl/+}$;Wnt1-Cre2* and *Hand2$^{fl/+}$* mice (N=6-8). **C|** B cells also increase from healthy to DSS-treated to AOM/DSS-treated colon for *Hand2$^{fl/+}$* mice, but are not increasing from inflamed to cancerous colon for *Hand2$^{fl/+}$;Wnt1-Cre2* mice. **D|** GC B cells only increase significantly in the cancer context in all mice, but the increase is dampened in the hypo-innervated mice.

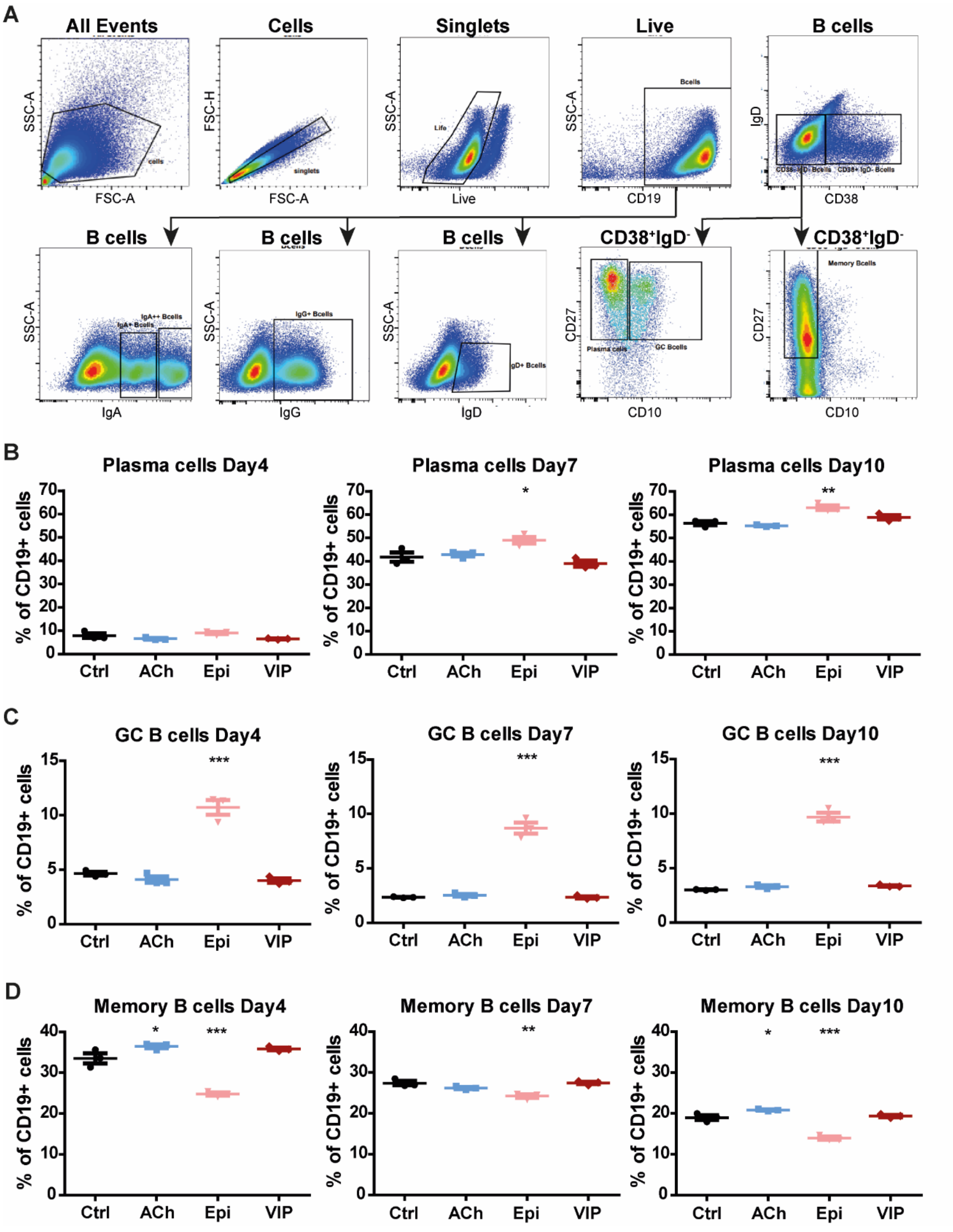


**Supplementary Figure 6| Gating strategy and flow cytometry analysis of human primary B cell cultures. A|** Flow cytometry analysis of primary human B cell cultures was performed on a Cytek® Aurora multispectral system using a B cell subtype panel. **B|** Plasma cells, characterized as CD38$^+$IgD$^-$CD27$^+$CD10$^-$, increase over time, but increase significantly more in the epinephrine-treated group (N=3) compared to control B cells (N=3) after 7 and 10 days of culture. **C|** GC B cells, identified as CD38$^+$IgD$^-$CD27$^+$CD10$^+$ cells, are not only increased after 4 days, but also after 7 and 10 days of epinephrine stimulation, while acetylcholine and VIP do not induce a response. **D|** CD38$^-$IgD$^-$CD27$^+$CD10$^+$ memory B cells slightly decrease over time, but are significantly lower at all timepoints than in control B cell cultures.

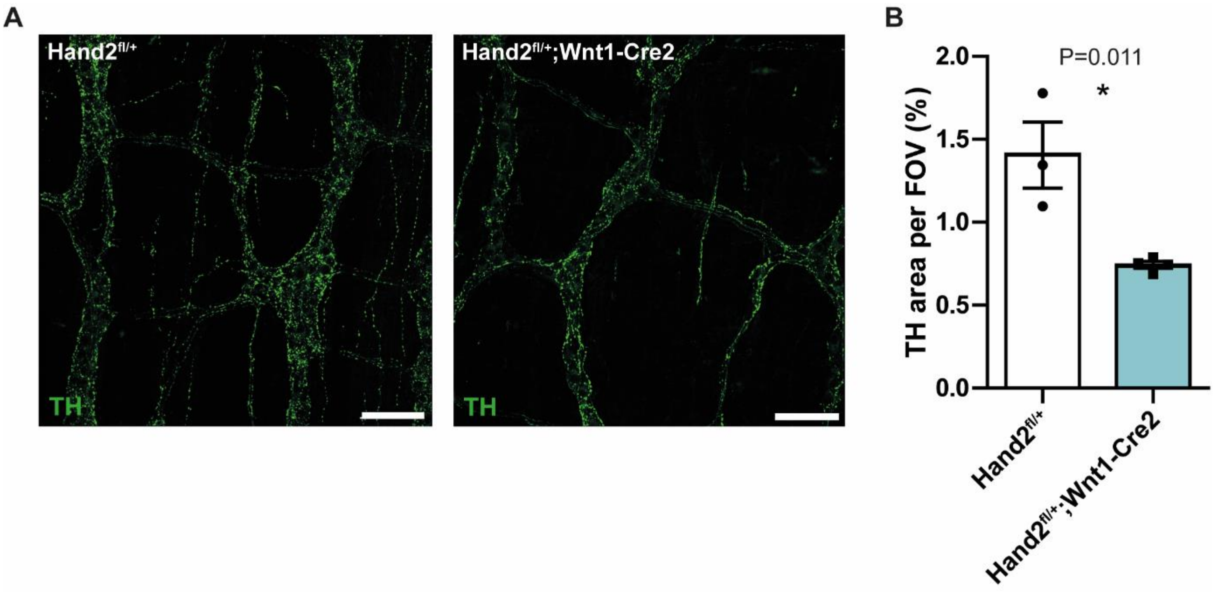


**Supplementary Figure 7| TH immunoreactivity in the hypo-innervated myenteric plexus of mouse colon.** A| Representative images of tyrosine hydroxylase (TH) immunofluorescent labelling represent B| a decrease in neuronal TH immunoreactivity in the myenteric plexus of colons from *Hand2*$^{fl/+}$*;Wnt1-Cre2* (N=4) compared to *Hand2*$^{fl/+}$ (N=3) mice. Scale bars are 100 µm.

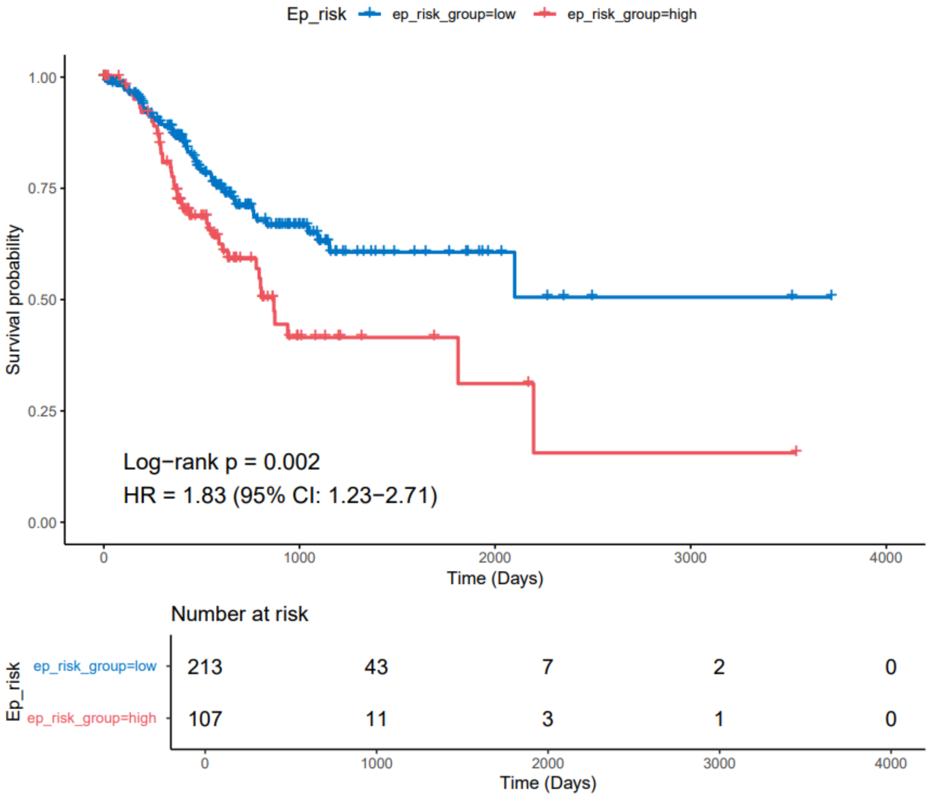


**Supplementary Figure 7| Association of the epinephrine-induced B cell signature with survival in the gastric cancer TCGA patient group.** The epinephrine-induced B cell signature (group high=red) is associated with a worse survival for gastric cancer patients as visualized with Kaplan-Meijer plot (P=0.002).
